# Glypican4 mediates Wnt transport between germ layers via signaling filopodia

**DOI:** 10.1101/2020.09.08.288613

**Authors:** Bo Hu, Anurag Kakkerla Balaraju, Juan J Rodriguez, Yuanyuan Gao, Nhan T Nguyen, Heston Steen, Saeb Suhaib, Songhai Chen, Fang Lin

**Author notes:** Corresponding author: Fang Lin.

## Abstract

Glypicans influence signaling pathways by regulating morphogen trafficking and reception. However, the underlying mechanisms in vertebrates are poorly understood. In zebrafish, Glypican 4 (Gpc4) is required for convergence and extension (C&E) of both the mesoderm and endoderm. Here we show that transgenic expression of GFP-Gpc4 in the endoderm of *gpc4* mutants rescues C&E defects in all germ layers. The rescue of mesoderm was likely mediated by Wnt5b and Wnt11f2, and depended on signaling filopodia rather than on cleavage of the Gpc4 GPI anchor. Gpc4 bound Wnt5b and regulated formation of the filopodia that transport Wnt5b to neighboring cells. Blocking signaling filopodia that extend from endodermal cells suppressed this rescue. Thus, endodermal signaling filopodia that expressed GFP-Gpc4 transported Wnt5b, and likely Wnt11f2, to other germ layers, rescuing the C&E defects caused by a *gpc4* deficiency. Our study reveals a new mechanism that could explain *in vivo* morphogen distribution involving Gpc4.

## Introduction

Glypicans (Gpcs), members of the heparan sulfate proteoglycan (HSPG) family, are bound to the external surface of the cell membrane by a C-terminal glycerophosphatidylinositide (GPI) anchor (Bulow and Hobert, 2006). Gpcs interact with numerous growth factors/morphogens such as Wnt, FGF, Bmp and hedgehog (Hh), thereby influencing a broad range of signaling pathways that regulate animal development (Poulain and Yost, 2015; Sarrazin, et al., 2011; Hacker, et al., 2005; Lin, 2004). Vertebrates have six Gpc isoforms (Gpc1-6) (De Cat and David, 2001). In humans, mutations in the *GPC3, GPC4* and *GPC6* genes are associated with congenital diseases such as Simpson-Golabi-Behmel (SGB) overgrowth syndrome (Campos-Xavier, et al., 2009; DeBaun, et al., 2001; Veugelers, et al., 1998; Pilia, et al., 1996). Studies in various animal models have revealed that Gpcs play roles in various developmental processes including: wing and eye development; the establishment of body length; jaw and heart development; and the migration of endodermal cells and cells of the lateral line primordium (LLP) (Hu, et al., 2018; Miles, et al., 2017; Venero Galanternik, et al., 2016; Strate, et al., 2015; Zhang, et al., 2013; LeClair, et al., 2009; Gumienny, et al., 2007). Thus, an understanding of Gpcs function will be important for understanding developmental processes as well as congenital defects and diseases.

Gpcs control extracellular signaling by various mechanisms. First, they act as co-receptors, stabilizing ligand-receptor interactions to enhance pathway activity (Yan, et al., 2010; Yan and Lin, 2007; Kan, et al., 1993). Second, Gpcs can act as repressors, either competing with morphogens for receptor binding (Capurro, et al., 2008) or recruiting a deacetylase to inhibit binding of a morphogen to its receptor (Kakugawa, et al., 2015). Third, Gpcs can control the diffusion or trafficking of morphogens to establish their gradients (Fico, et al., 2011; Filmus, et al., 2008). They do so by several mechanisms: shedding of Gpcs from the cell surface via cleavage at the GPI anchor (Kreuger, et al., 2004), which can change the concentrationof a morphogen in the local environment as well as its availability to adjacent cells; induction of endocytosis and subsequent degradation to remove morphogens from the cell surface (Capurro, et al., 2008); internalization of a morphogen to spread it to neighboring cells via transcytosis (Callejo, et al., 2011; Gallet, et al., 2008); the recruitment of lipophorin vesicles to transport and release morphogens to distant cells (Eugster, et al., 2007; Panakova, et al., 2005); and expression in migrating cells to deliver morphogens to the distant locations (Serralbo and Marcelle, 2014).

Morphogens can be moved not only by trafficking, but also by actin-based signaling filopodia, known as cytonemes, which can transport morphogens in distance to their sites of action (Gonzalez-Mendez, et al., 2019; Ramirez-Weber and Kornberg, 1999). Notably, in the zebrafish blastula, formation of the signaling filopodia that transport Wnt8a can be induced by non-canonical Wnt/planar cell polarity (Wnt/PCP) signaling (Mattes, et al., 2018). Additionally, *Drosophila* Gpcs, Dally and Dlp are shown to coat cytonemes that transport Hh (Gonzalez-Mendez, et al., 2017), suggesting that Gpcs play a role in cytoneme formation. Although many mechanisms have been proposed to explain the actions of HSPGs in the distribution and function of morphogens, the mechanisms whereby Gpcs regulate morphogens in vertebrates remain poorly understood.

In zebrafish and *Xenopus*, Gpc4 was first indentified as a positive modulator of Wnt11f2 in regulating mesodermal convergence and extension (C&E) (Ohkawara, et al., 2003; Topczewski, et al., 2001), a process that narrows and elongates all three germ layers (ectoderm, mesoderm, and endoderm) to establish the animal body plan (Solnica-Krezel and Sepich, 2012; Keller, 2002). Later studies showed that Gpc4 is versatile, contributing to many additional developmental processes via its influence on signaling by several pathways (Shh, BMP, and Wnt signaling) (Miles, et al., 2017; Venero Galanternik, et al., 2016; Strate, et al., 2015; LeClair, et al., 2009). However, little is known about how Gpc4 affects morphogens *in vivo*.

Recently, we and others showed that Gpc4 is required for endoderm C&E, in both the anterior and posterior regions (Hu, et al., 2018; Miles, et al., 2017). To investigate the cell autonomy of Gpc4 in the gut endoderm, we generated a transgenic line that expresses Gpc4 specifically in this germ layer. Intriguingly, in *gpc4*^*-/-*^ embryos the endodermal expression of GFP-Gpc4 partially, but significantly, rescued the C&E defects in all germ layers, suggesting that Gpc4 functions both within and outside the endoderm. Thus, our animal model provides a unique opportunity to explore the mechanisms underlying communication among germ layers. We found that although GFP-Gpc4 could be cleaved and delivered, the rescue was not due to Gpc4 cleavage at the GPI anchor. Instead, GFP-Gpc4 present in the signaling filopodia of endodermal cells that transport Wnt5b to neighboring cells was responsible. Thus, our study uncovers a new mechanism, whereby the contribution of Gpc4 to the formation of signaling filopodia accounts for its non-cell autonomous functions.

## Results

### Endodermal expression of Gpc4 rescues C&E defects in all germ layers of *gpc4*^*-/-*^ embryos

Previously, we detected *gpc4* transcript in the anterior endoderm (Hu, et al., 2018). Here, we found that *gpc4* is also expressed in the posterior endoderm (Fig. S1A-C′′), from which the gut will develop. To evaluate gut formation in *gpc4* mutants, we assessed the expression of *foxa3* by whole-mount *in situ* hybridization (WISH) (Mayer and Fishman, 2003). Compared to the gut-tube in control (sibling) embryos, that in *gpc4*^*-/-*^ embryos was significantly enlarged and the digestive organs were malformed (duplicated, smaller, or missing; Fig. 1B vs A, and data not shown). To determine whether Gpc4 regulates morphogenesis of the gut endoderm cell-autonomously, we generated transgenic line *Tg*(*sox17:GFP-gpc4*), in which expression of GFP-tagged Gpc4 is driven by the endoderm-specific promoter *sox17* (Woo, et al., 2012). GFP was inserted immediately after the N-terminal signal peptide of Gpc4 to avoid disrupting its membrane localization (Fig. S1D). GFP-Gpc4 is functional, as injection of the encoding RNA rescued C&E defects in *gpc4* mutants (Hu, et al., 2018). Consistent with the expression pattern of *sox17* (Alexander, et al., 1999), in this line GFP-Gpc4 signal was detected in the innermost tissue of the embryo, including in Kupffer’s vesicle at the tailbud (TB) stage (Fig. S1E-E′′). Additionally, when *Tg*(*sox17:GFP-gpc4*) zebrafish were outcrossed to *Tg*(*sox17:memCherry*) counterparts, in which mCherry is expressed in the plasma membrane of endodermal cells (Ye, et al., 2015), GFP-Gpc4 co-localized with mCherry (Fig. S1F-F′′). For the studies described here, we used a line in which expression of GFP-Gpc4 was modest and embryogenesis was normal.

**Figure 1.**
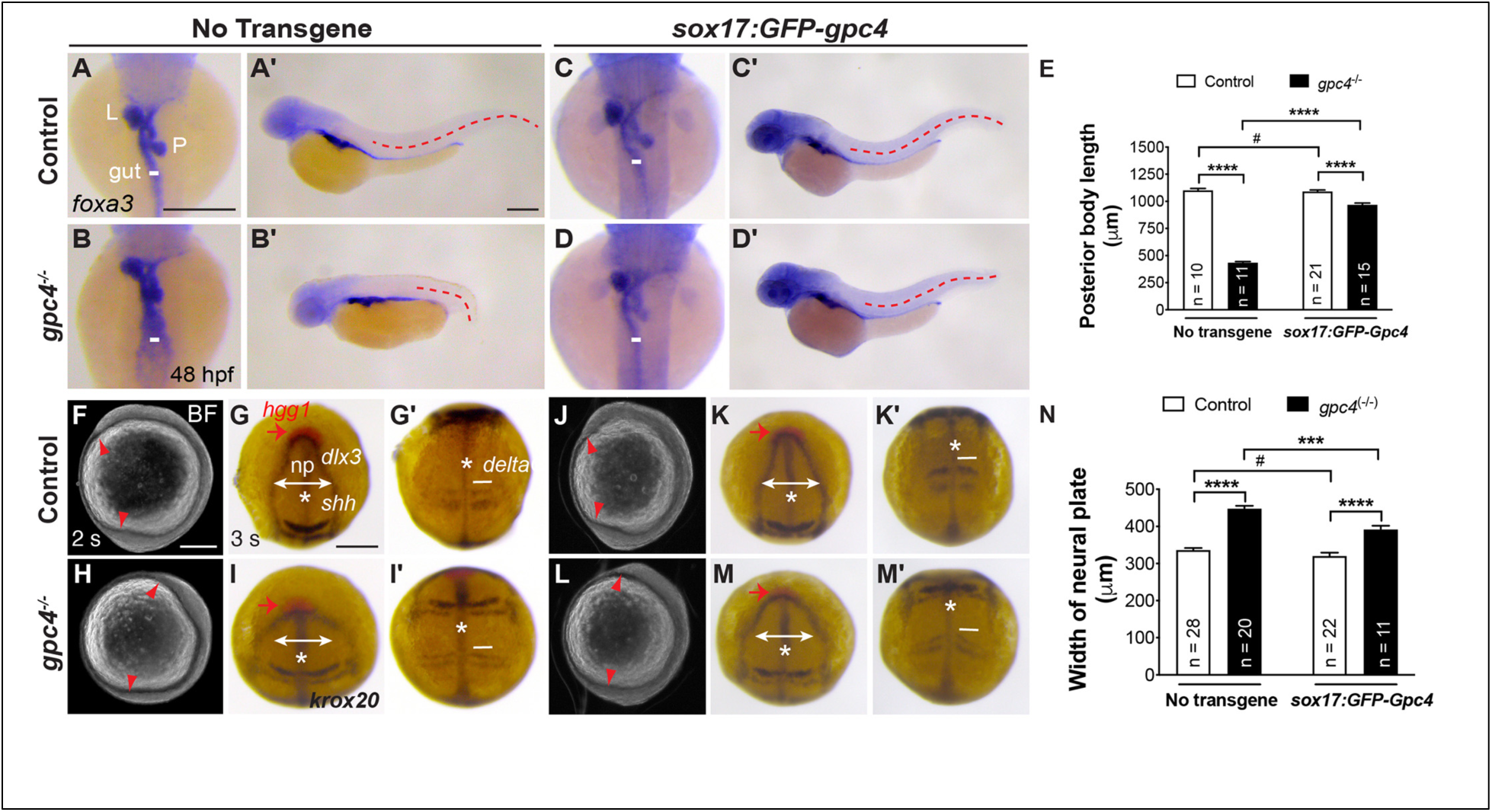
Endodermal expression of GFP-Gpc4 rescues C&E defects in all germ layers of *gpc4*^*-/-*^ embryos. (A-D′) The expression of *foxa3* as detected by whole-mount *in situ* hybridization (WISH) of the indicated embryos at 48 hpf, showing the morphology of the gut, liver (L), and pancreas (P). (A-D) Dorsal view. (A′-D′) Lateral view. White lines, width of the gut-tube, lines in all embryos are equal in length; Red dashed lines, posterior body length. (E) Average posterior body length in embryos shown in (A′-D′). (F, H, J, L) Bright-field images of the indicated embryos at 2s. Red arrowheads point to the anterior and posterior points of the embryonic axis. Lateral view. (G, I, K, M) Expression of *hgg1* (red), *dlx3, krox20, shh, and deltaC* at 3s, as detected by WISH. *, notochord (*shh*); np, neural plate (*dlx3*); white lines with double arrows, width of the neural plate; white lines (*deltaC*), width of the first somite. All lines of each type are equal in length. (J-M) Dorsoanterior view; (J′-M′) Dorsal view. (N) Average width of neural plate in embryos shown in (J-M). # P>0.05, ***P<0.001, ****P<0.0001; Student’s t-test. Scale bars: 200 μm.

To determine whether endodermal defects in *gpc4*^*-/-*^ embryos are due to *gpc4* deficiency specifically in the endoderm, we generated the *gpc4*^+*/-*^/*Tg*(*sox17:GFP-gpc4*) line. Strikingly, the *gpc4*^*-/-*^/*Tg*(*sox17:GFP-gpc4*) offspring derived from incrossing this line did not exhibit the short body axis that is typical for *gpc4*^*-/-*^ embryos (Fig. 1D′ vs B′). To evaluate mesodermal C&E defects, we assessed the length of the posterior body. Expression of GFP-Gpc4 in the endoderm did not affect posterior body length in controls, but it largely suppressed the shortening of posterior body length in the *gpc4*^*-/-*^ embryos (Fig.1A′-D′, E), indicating that endodermal expression of GFP-Gpc4 rescues mesodermal C&E defects. Notably, the body length in the *gpc4*^*-/-*^*/Tg*(*sox17:GFP-gpc4*) embryos was shorter than that in control embryos (Fig. 1D′ vs C′), suggesting that the rescue was not complete. Additionally, the morphology of gut-tube and digestive organs in *Tg*(*sox17:GFP-gpc4*) embryos is normal (Fig. 1C), suggesting that expressing GFP-Gpc4 in the endoderm in this line does not affect normal development of the digestive system. Notably, in *gpc4*^*-/-*^*/Tg*(*sox17:GFP-gpc4*) embryos, the enlargement of the gut tube and malformation of organs associated with the *gpc4*^*-/-*^ genotype were absent (Fig. 1D vs B). These data are consistent with endodermal expression of Gpc4 rescuing the C&E defects of both the mesoderm and endoderm in *gpc4* mutants.

To determine when such rescue occurs, we examined the embryos during early segmentation, at which point mesodermal C&E defects in *gpc4*^*-/-*^ embryos are prominent (Topczewski, et al., 2001). At the 2 somite stage (2s), the body axes of *gpc4*^*-/-*^ */Tg*(*sox17:GFP-gpc4*) embryos were significant elongated relative to those of *gpc4*^*-/-*^ embryos (Fig. 1L vs H). Whole-mount *in situ* hybridization (WISH) for tissue-specific markers of the neural plate (*dlx3*), notochord (*shh*), prechordal plate cells (*hgg1*), somites (*deltaC*), rhombomeres 3 and 5 (*krox20*, for the purpose of staging) in 3s-embryos revealed a significant suppression of the broadening of the neural plate, notochord, and somites typically seen in *gpc4*^*-/-*^ embryos (Fig. 1M,M′ vs 1I,I′, N). Thus, the resecue of mesoderm and ectorderm defects by endodermal expression of GFP-Gpc4 was evident at early segmentation, suggesting that the rescue effect is long-range and that Gpc4 functions both within and outside of the endoderm.

### The rescue of C&E defects in *gpc4* mutants by endodermal expression of GFP-Gpc4 is dependent on the endoderm

To exclude the possibility that the *sox17* promoter induces expression of GFP-Gpc4 outside the endoderm, which could contribute to the observed rescue, we employed two approaches that eliminate formation of the endoderm. If the observed rescue indeed resulted from endodermal expression of Gpc4, removal of the endoderm should abolish the rescue. First, we eliminated this germ layer by injecting embryos with a morpholino (MO) targeting *sox32*, a transcription factor required for development of the endoderm (Dickmeis, et al., 2001). Injection of the *sox32* MO did not affect posterior body length in either the control or *gpc4*^*-/-*^ embryos; as expected, it produced pericardial edema in both because endoderm deficiency disrupts heart development (Fig. S2E vs B and not shown) (Alexander, et al., 1999). Notably, *gpc4*^*-/-*^*/Tg*(*sox17:GFP-gpc4*) embryos injected with this MO were short and similar to *gpc4*^*-/-*^ embryos (Fig. S2F vs B), indicating that the *sox32* MO largely suppressed the rescue of the body length (Fig. S2F vs D). Second, we generated *gpc4/sox32* double mutants. As was the case for MO-injected embryos, the posterior body length of *sox32*^*-/-*^ embryos was comparable to that of the sibling control embryos, and *sox32* deficiency did not affect this parameter in *gpc4*^*-/-*^ embryos (Fig. 2A-D, I). The *sox32* deficiency did not affect posterior body length in *Tg*(*sox17:GFP-gpc4*) embryos (Fig. 2F vs E, J), yet it led to significant shortening in *gpc4*^*-/-*^ embryos (Fig. 2H vs G, J). Thus, such rescue of the body axis is dependent on the endoderm.

**Figure 2.**
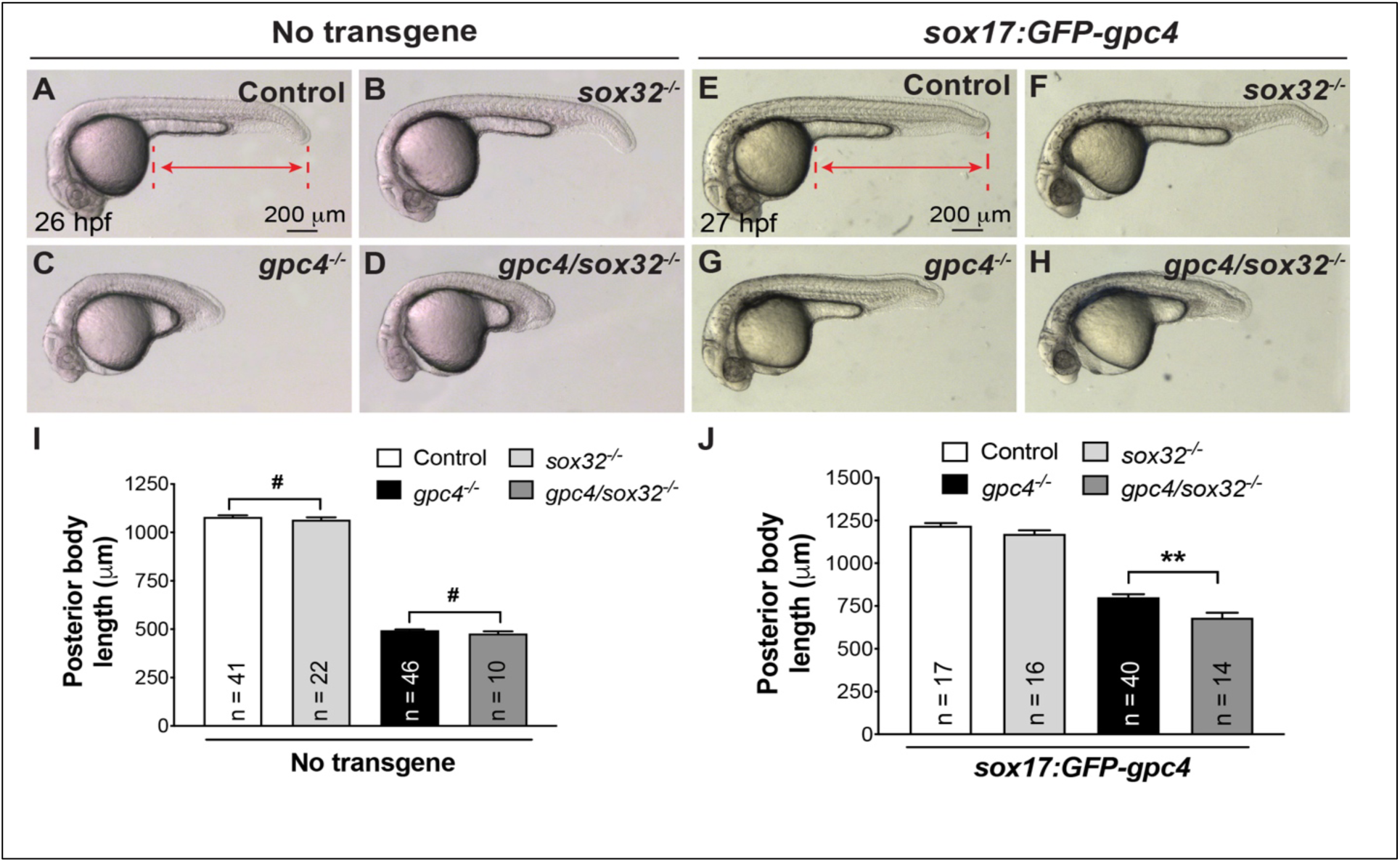
The rescue of C&E defects in *gpc4* mutants by endodermal expression of GFP-Gpc4 is dependent on the endoderm. (A-H) Bright-field images of the indicated embryos. Lateral view. Red lines with double arrows, posterior body length; dashed lines, points used to measure posterior body length. (I) Posterior body length in embryos shown in (A-D). (J) Posterior body length in embryos shown in (E-H). The number of embryos tested in each group is indicated on each bar. # P>0.05, **P<0.01; Student’s t-test.

### The rescue of mesodermal C&E by endodermal expression of GFP-Gpc4 is partially mediated by Wnt5b and Wnt11f2

As Wnt/PCP signaling is the major means of regulating mesodermal C&E (Solnica-Krezel & Sepich, 2012; Tada, Concha, & Heisenberg, 2002), we postulated that endodermal Gpc4 mediates the rescue of mesodermal C&E by influencing Wnt/PCP signaling. In zebrafish, both Wnt5b and Wnt11f2 are involved in Wnt/PCP signaling (Kilian, et al., 2003; Heisenberg, et al., 2000) and a previous study had shown that *Xenopus* Gpc4 can physically bind Wnt5, Wnt8 and Wnt11 (Ohkawara, et al., 2003). We further examined the ability of zebrafish Gpc4 to bind Wnt5b and Wnt11f2. To this end we transfected HEK293 cells with Flag-Gpc4 or Flag-JNK (as a negative control), and myc-Wnt5b or myc-Wnt11f2, and performed co-immunoprecipitation of cell lysates. Flag-Gpc4, but not Flag-JNK, was pulled down with myc-Wnt5b or myc-Wnt11f2 (Fig. 3A, B). Thus, zebrafish Gpc4 interacts physically with Wnt5b and Wnt11f2. Notably, our pull-down assay showed that Flag-Gpc4ΔGAG, a Gpc4 mutant lacking the HS chain-binding domain of Gpc4 (Fig. 4A and B), binds Wnt5b and Wnt11f2 (Fig. 3A and B, Lane 2), suggesting that the ability of zebrafish Gpc4 to bind Wnt proteins is not dependent on the HS chains attached to its GAG domain.

**Figure 3.**
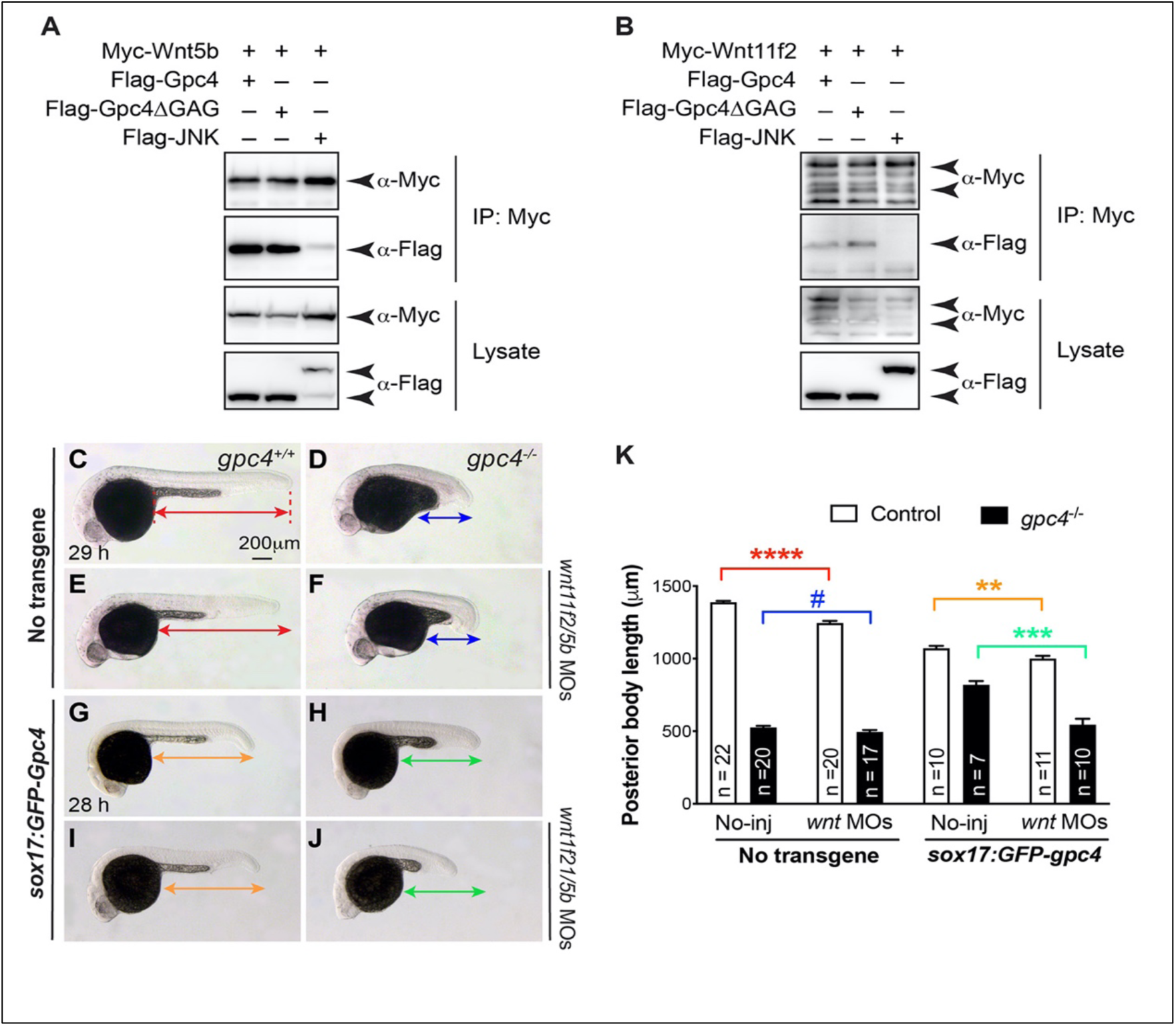
The rescue of mesodermal C&E by endodermal expression of GFP-Gpc4 is mediated in part by Wnt5b and Wnt11f2. (A, B) Western blots of co-IP experiments with Myc-tagged Wnt5b (E) or Wnt11f2 (F) and Flag-tagged full-length Gpc4, Gpc4ΔGAG, or JNK (negative control) in both pellets (IP) and cell lysates (Lysate). Blotting was performed using anti-Flag and anti-Myc antibodies. (C-J) Bright-field images of the indicated control embryos and embryos injected with a sub-dose of MOs targeting *wnt11f2/5b* (5 and 1ng). Lateral view. Lines with double arrows, posterior body length. Lines of the same color are equal in length. (K) Average posterior body length in embryos shown in (A-H). The number of embryos analyzed in each group is indicated in each bar. P values in different colors correspond to the embryos with posterior body length marked by line of that color. #, p>0.05; **, P<0.01; ***, P<0.001; ****, P<0.0001; Student’s t-test.

**Figure 4.**
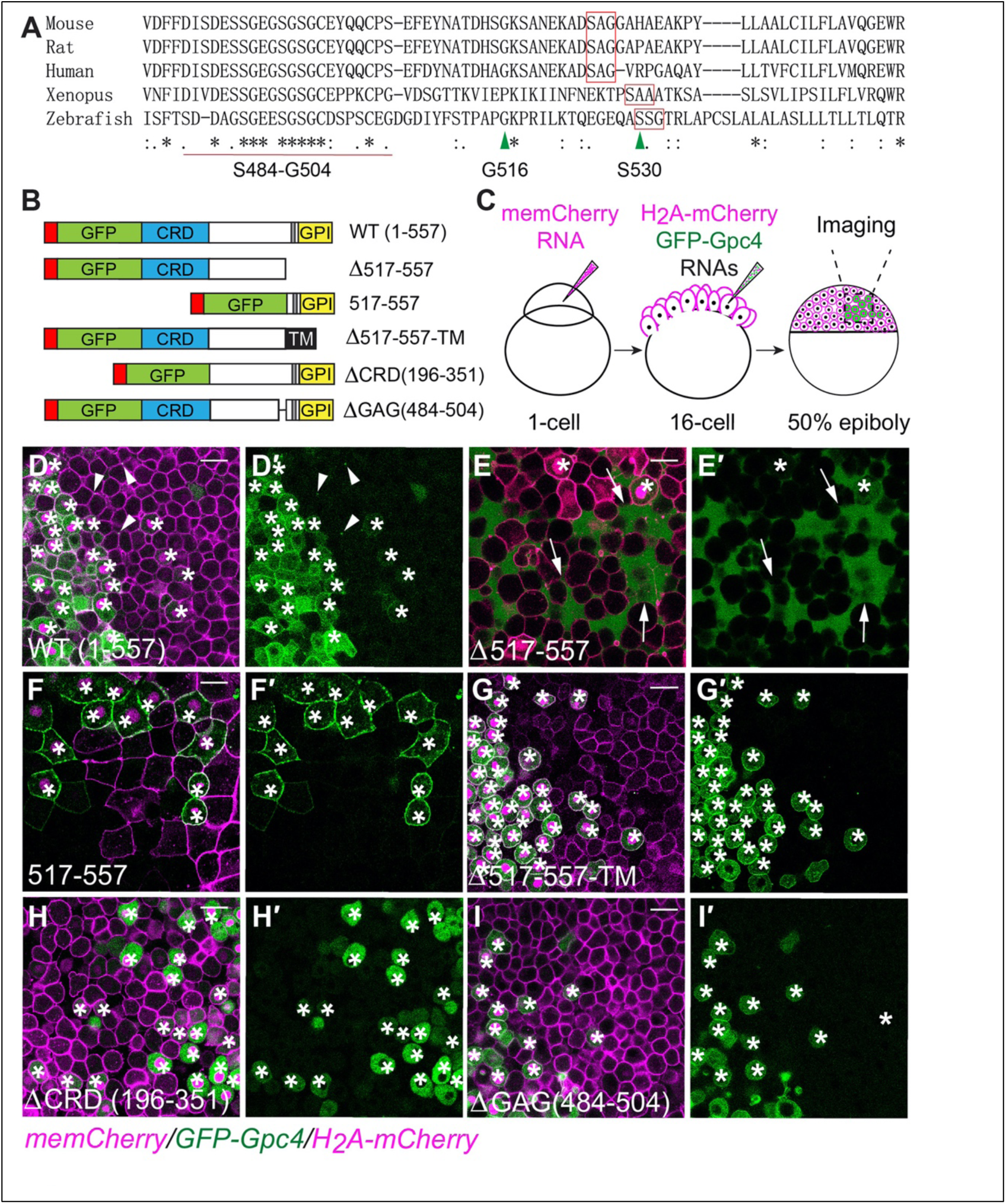
GFP-Gpc4 mutant proteins localize to distinct sites in the zebrafish blastula. (A) Alignment of C-terminal amino acids of Gpc4 from the indicated species. Stars (*), identical amino acids; red rectangles, putative GPI attachment sites; red line, conserved HS chain attachment domain (GAG domain, S484-G504); first green arrowhead (S530), putative GPI attachment site in zebrafish; second green arrowhead (G516), site of fusion to the transmembrane domain (TM). (B) Schematic of WT, truncation, deletion, and fusion construct forms of Gpc4. Red rectangles indicate N-terminal signal peptide. Gray rectangles indicate putative GPI attachment sites. (C) Schematic diagram illustrating the *in vivo* localization assay. (D-I′) Confocal images of blastulas, with all cells labeled with memCherry (in megenta) and some cells co-labeled with the indicated GFP-Gpc4 constructs and nuclear H_2_A-mCherry (*). White arrowheads (D-D′), punctate GFP signal on membrane; white arrows (E-E′), the extracellular space outside GFP-Gpc4-expressing cells (E-E′). Scale bars: 20 μm

To determine whether endodermal GFP-Gpc4 is likely to regulate mesoderm C&E by influencing the expression of Wnt5b and Wnt11f2, we tested the involvement of Wnt5b and Wnt11f2 in morphogenesis of the mesoderm and endoderm at 48 hpf. The posterior body length and *foxa3* expression patterns were tested as described in Fig. 1 and the posterior body length of *wnt11f2*^*-/-*^ embryos was found to be similar to that of control siblings, but that of *wnt5b*^*-/-*^ embryos was significantly shorter and that of double *wnt11f2*^*-/-*^*/wnt5b*^*-/-*^ embryos was shortest (Fig. S3A-D, E). Phenotypes related to endoderm morphology were consistent with these findings: the gut-tube was normal in *wnt11f2*^*-/-*^ embryos but slightly enlarged in *wnt5b*^*-/-*^ and significantly widened in *wnt11f2*^*-/-*^*/wnt5b*^*-/-*^ mutants (Fig. S3A′-D′). These data suggest that at day 2, Wnt5b but not Wnt11f2 is required for elongation of the body axis and for formation of the gut-tube, but Wnt11f2 co-operates Wnt5b to regulate endoderm morphogenesis.

Given the observed rescue by endodermal GFP-Gpc4 as early as 2s (Fig. 1), we examined the expression of tissue markers of C&E at 3s in the Wnt mutants. Consistent with published data (Heisenberg, et al., 2000), we found that in *wnt11f2*^*-/-*^ embryos, cells of the prechordal plate (*hgg*) failed to migrate as far anterior to the neural plate (*dlx3*, np) as their control counterparts that and the neural plate and notochord (*shh*) were broader (Fig. S3G-G′ vs F-F′, J). However, in the anterior region of *wnt5b*^*-/-*^ embryos, the positions of the prechordal plate cells and the widths of the notochord and neural plate appeared to be normal, whereas in the posterior region the neural plate was slightly wider (Fig. S3H-H′, J). In *wnt11f2/wnt5b* double mutants, defects in both the mesoderm and ectoderm were much more severe that those in the single mutants (Fig. S3I-I′, J), consistent with a previous report on effects at the tailbud stage (Kilian, et al., 2003). These data indicate that during early segmentation, *wnt11f2* is required for C&E of both the mesoderm and ectoderm, in the anterior as well as the posterior region, and that *wnt5b* affects ectodermal C&E only in the posterior region.

Considering the relatively normal body length in *wnt11f2*^*-/-*^ embryos and the shorter body length in *wnt5b*^*-/-*^ embryos at day 2, it appears that *wnt5b* functions at a later stage of embryogenesis (segmentation) than wnt11f2 (during gastrulation). However, consistent with what we found at day 2, *wnt5b* and *wnt11f2* functioned synergistically in regulating the C&E of all germ layers at early segmentation. Therefore, both Wnt5b and Wnt11f2 might contribute to endodermal, GFP-Gpc4-mediated rescue of mesoderm C&E defects in *gcp4* mutants.

To test this possibility, we tested whether loss of *wnt5b* and *wnt11f2* can suppress the rescue of C&E defects by endodermal expression of Gpc4. Given that *wnt5b/wnt11f2* double mutants have C&E defects, it was not possible to use them for this analysis. We reasoned that if the rescue of Gpc4 depends on these Wnt proteins, embryos should be more sensitive to the suppression of their expression. Thus, we injected embryos with MOs targeting both *wnt5b* and *wnt11f2* at sub-doses that partially suppress their expression. Injecting MOs at these doses led to a slight but significant reduction in posterior body length in control embryos, regardless of whether GFP-Gpc4 was expressed in the endoderm (Fig. 3E vs C, I vs G, K). Intriguingly, although such MO injection did not affect the body length in *gpc4*^*-/-*^ embryos (Fig. 3F vs D, K), it suppressed the rescue caused by endodermal GFP-Gpc4. In embryos injected with MOs, posterior body length was significantly shorter than that in the uninjected controls (Fig. 3J vs H, K). These results indicate that in *gpc4*^*-/-*^ embryos the rescue of mesodermal C&E defects induced by endodermal expression of GFP-Gpc4 is at least partially mediated by Wnt5b and Wnt11f2.

### The ability of endodermal GFP-Gpc4 to rescue body length is not dependent on the GPI anchor

We set out to identify the mechanisms whereby endodermal expression of GFP-Gpc4 exerts a long-range rescue effect on other germ layers. One feature of the GPI-anchored proteins (GPI-APs) is that these anchors can be cleaved by various enzymes to influence GPI-AP function (Fujihara and Ikawa, 2016). For example, Notum, an *α*/*β*-hydrolase can cleave *Drosophila* glypicans Dally-like and Dally (Kreuger, et al., 2004), as well as mammalian GPC3, GPC5, and GPC6 (Traister, et al., 2008). However, zebrafish Notum does not interact with and cleave zebrafish Gpc4 (Flowers, et al., 2012). Nonetheless, when GPI biosynthesis is disrupted, zebrafish Gpc4 localization shifts from the membrane to the extracellular space, suggesting that the GPI anchor is critical for its attachment to the cell membrane (Shao, et al., 2009). Thus, it is possible that GFP-Gpc4 expressed on the endoderm can be cleaved and released into the extracellular space, and thus that it can function outside the endoderm, in the mesoderm, and/or in the ectoderm.

To test this hypothesis, we first used a blastula assay to assess GFP-Gpc4 for cleavage *in vivo* (Fig. 4C). Briefly, we injected embryos at the one-cell stage with the memCherry RNA to label the plasma membrane of all cells. When these embryos reached the 16-cell stage, one blastula was co-injected with RNAs encoding *H*_*2*_*A-mCherry* and *GFP-gpc4* to label a subset of cells, with the nuclei of GFP-Gpc4 expressing cells labeled with mCherry. Live imaging of embryos at 50% epiboly showed that GFP-Gpc4 was expressed in both the cytosol and the plasma membrane of the cells whose nuclei express mCherry; however, it was also detected in neighbors several cells away from the site of expression, on the membrane and sometimes as bright puncta at sites of cell-cell contact (Fig. 4D, D′). These data suggest that GFP-Gpc4 can be delivered to cells near those in which it is expressed.

Next, we sought to identify the GPI region of zebrafish Gpc4. GPI is synthesized in the endoplasmic reticulum (ER) and transferred to the attachment site within the C-terminal region of Gpcs (Kinoshita and Fujita, 2016; Watanabe, et al., 1995). We analyzed the amino acid sequences in the C-terminal regions of Gpc4 from a variety of species, including mouse, rat, human, *Xenopus* and zebrafish. Compared with its orthologs from other species, zebrafish Gpc4 is not well conserved in this region. We nevertheless identified putative conserved GPI attachment sites: SSG in zebrafish, SAG in mammals, and SAA in *Xenopus* (Fig. 4A). We generated a series of C-terminal truncation mutants (Fig. 4B) and assessed their expression patterns using the *in vivo* blastula assay, as described above. Gpc4 lacking AA517-557 (GFP-Gpc4Δ517-557) failed to localize to the cell membrane. Instead, it was present mainly in the extracellular space (Fig. 4E, E′). In contrast, GFP-fused Gpc4AA517-557 was expressed mainly on the cell membrane (Fig. 4F, F′). Thus, AA517-557 of Gpc4 encompasses the GPI attachment signal peptide, which is critical for the membrane anchor.

We postulated that if the rescue stems from release of Gpc4 after GPI cleavage, a mutated Gpc4 that cannot be cleaved should not be able to rescue the mesodermal defects. Thus, we generated a chimeric construct, GFP-Gpc4Δ517-557-Sdc4TM, in which the GPI anchor was replaced with the transmembrane domain (TM) of Syndecan 4 (Sdc4), another HSPG family member that binds the cell membrane, but in this case via its single TM domain rather than a GPI anchor (Lopes, et al., 2006; Munoz, et al., 2006). *In vivo* localization revealed that GFP-Gpc4Δ517-557-Sdc4TM was expressed on the cell membrane (Fig. 4G, G′). Thus, the TM domain restores membrane localization to GFP-Gpc4 lacking the GPI anchor. Furthermore, this chimeric construct was functional, as its overexpression rescued the shortened body length in *gpc4* mutants (Fig. S4A-D). We next generated transgenic line *Tg*(*sox17:GFP-gpc4*Δ*517-557-sdc4TM*), in which to express the chimeric protein specifically in the endoderm. A stable transgenic line that exhibited normal embryogenesis was crossed to the *gpc4*^+*/-*^ zebrafish. Like *gpc4*^*-/-*^*/Tg*(*sox17:GFP-gpc4*) embryos, *gpc4*^*-/-*^*/Tg*(*sox17:GFP-gpc4*Δ*517-557-sdc4TM*) embryos had a longer body length and gut-tube than their *gpc4*^*-/-*^ counterparts, and normal digestive organs (Fig. S4F, F′ vs E, E′). Thus, endodermal expression of the chimeric protein that lacks the cleavable GPI signal but retains membrane localization was able to rescue the C&E defects in both the mesoderm and endoderm of *gpc4*^*-/-*^ embryos, indicating that GPI cleavage of Gpc4 does not drive rescue of the mesoderm.

### Gpc4 contributes to the formation of actin-based signaling filopodia that transport Wnt5b

Recently, it was shown that some morphogens, including Wnt, can be transported to cells by specialized cell protrusions called signaling filopodia, actin-based structures that bind to and transport signaling molecules, enabling them to function at a distance from their site of expression (Stanganello and Scholpp, 2016; Jacquemet, et al., 2015; Kornberg and Roy, 2014). In our blastula assay, we mosaically labeled a subset of zebrafish blastula cells (whose plasma membranes were labeled with mCherry) with GFP-Gpc4 and nuclear H_2_A-mCherry. We also detected GFP-Gpc4-expressing cells that extend GFP-positive protrusions (Fig. S5A-A′′). Similarly, we observed that in *Tg*(*sox17:GFP-gpc4*) embryos, GFP-Gpc4-expressing endodermal cells extended robust GFP-positive protrusions (Movie 1). To determine whether GFP-Gpc4-positive protrusions are also actin-based, we performed mosaic injection of embryos with RNAs encoding *GFP-gpc4* and *RFP-Lifeact* (an F-actin binding protein that marks filopodia) (Mattila and Lappalainen, 2008; Riedl, et al., 2008). Live imaging showed that Lifeact-RFP illuminates GFP-Gpc4-labeled filopodia (Fig. S5B-B′′), suggesting that Gpc4 contributes to the formation of actin-based filopodia.

Our co-IP experiments showed that Gpc4 can physically bind Wnt5b, suggesting that GFP-Gpc4-labeled filopodia from endodermal cells can bind to and transport Wnt5b. To test this possibility, we injected *Tg*(*sox17:GFP-gpc4*) embryos with an *mCherry-wnt5b* RNA and performed confocal time-lapse experiments on the endoderm in the embryo posterior. Indeed, mCherry-Wnt5b puncta were present at GFP-Gpc4-labeling filopodia (Fig 5A,B). Notably, two types of Wnt5b-positive protrusions were observed: one extended to deliver Wnt5b out of the cells (Fig. 5A, yellow arrowheads), and the other retracted to carry Wnt5b back to cells (Fig. 5A, white arrowheads). In some cases, Wnt5b-positive protrusions from two adjacent cells contacted each other (Fig. 5A, cyan arrowheads). We also observed that Wnt5b-positive protrusions from distant cells formed a bridge connecting them (Fig. 5B, cyan arrowheads), indicating that that GFP-Gpc4-labeled filopodia can transport Wnt5b to other cells. To determine the role of Gpc4 in the formation of protrusions in endodermal cells, we performed confocal time-lapse of *Tg*(*sox17:memGFP*) embryos, in which all protrusions are labeled with GFP. We found that endodermal cells extended robust finger-like protrusions (Movie 2) and that: some of these extended to the space between endodermal cells (Fig. S6A and B, white arrowheads); some reached the neighboring endodermal cells (Fig. S6A and B, yellow arrowheads); and some contacted and connected with protrusions emerging from other endodermal cells (Fig. S6A and B, cyan arrowheads). These findings suggest that these filopodia communicate with other cells. Notably, in *gpc4* mutant embryos, endodermal cells formed shorter protrusions that might not be able to reach other distant cells (Fig. S6B). Quantification revealed that in *gpc4*^*-/-*^ embryos the total number of protrusions was comparable to that in controls, but the average length of the protrusions was shorter and the portion of long protrusions (>7 μm) was significantly lower (Fig. S6C-E). These data suggest that Gpc4 is critical for generating long protrusions that enable communication between distant cells.

**Figure 5.**
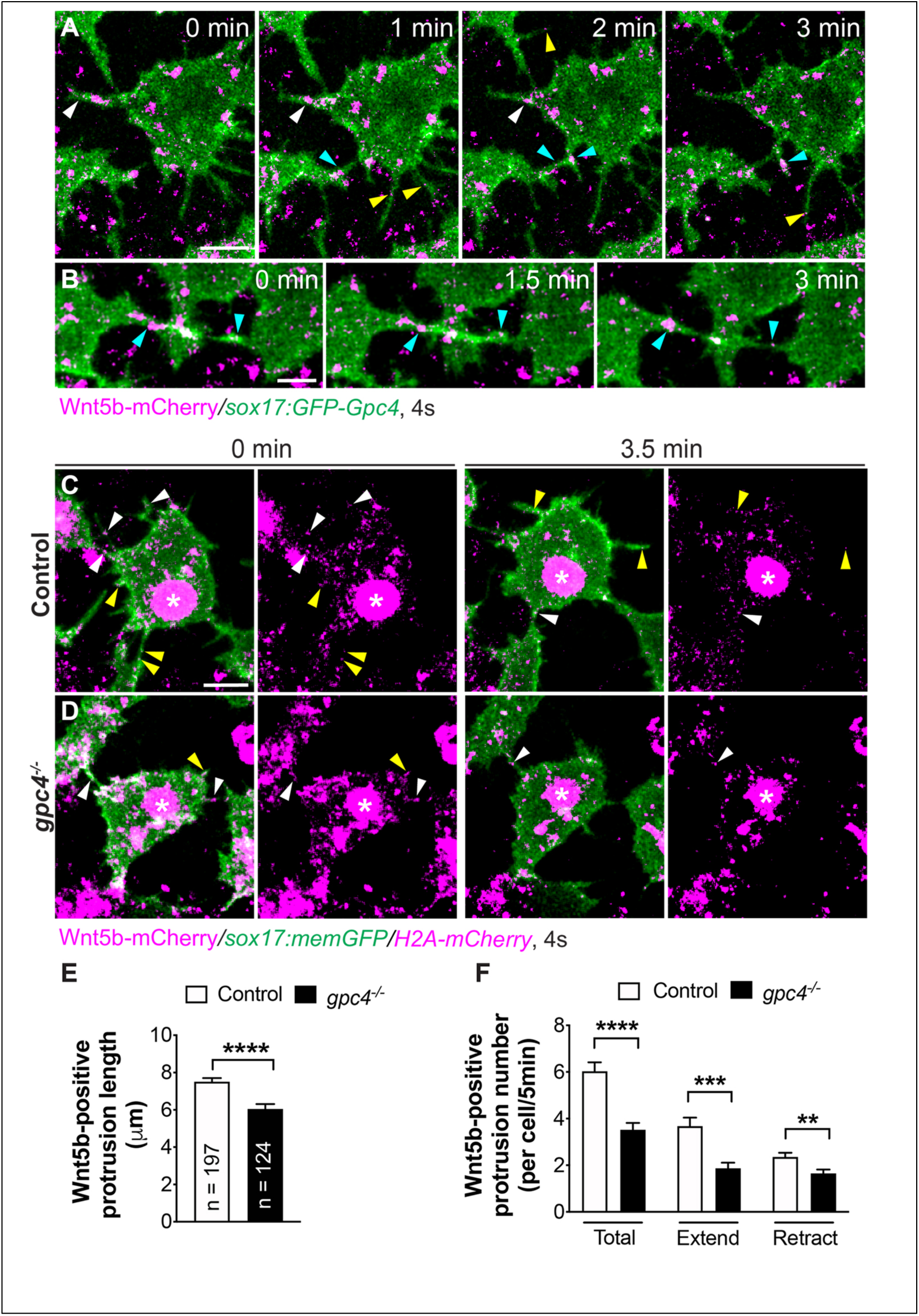
Gpc4 is required for the formation of Wnt5b positive filopodia. (A-B) Snapshots from confocal time-lapse images of *Tg*(*sox17:GFP-gpc4*) embryos injected with *wnt5b*-*mCherry* synthetic RNA. Yellow arrowheads, protrusions extending out; white arrowheads, protrusions retracting back; cyan arrowheads, protrusions merging from two cells connected. (C-D) Snapshots from confocal time-lapse images of *Tg*(*sox17:memGFP*) embryos injected with *wnt5b*-*mCherry* synthetic RNA, showing that Wnt5b-positive filopodia are present in both control and *gpc4*^*-/-*^ embryos. Wnt5b-mCherry (in magenta) is present on extending filopodia (yellow arrowheads), retracting filopodia (white arrowheads), and connecting filopodia emerged from two cells (cyan arrowheads). (E-F) Average length of protrusions (E) and frequency (F) of formation of Wnt5b-positive filopodia per endodermal cell during a 5-min window, in control embryos (197 protrusions, 19 cells, 4 embryos) and *gpc4*^*-/-*^ embryos (124 protrusions, 19 cells, 4 embryos). **, P<0.01; ***, P<0.001; ****, P<0.0001; Student’s t-test. Scale bars: 10 μm.

Next, we determined how Gpc4 affects Wnt5b transport. We injected embryos obtained from incrossing *gpc4*^+*/-*^/*Tg*(*sox17:memGFP*) fish with *wnt5b-mcherry* RNA, and performed confocal time-lapse. Wnt5b-Cherry labeled signaling protrusions extending from endodermal cells were examined. As we had observed in *Tg*(*sox17:GFP-gpc4*) embryos, protrusions extended to and retracted from cells (Fig. 5C and D). In *gpc4* mutant embryos, we observed that not only protrusions were shortened like those in Fig. S6 (Fig. 5E), but also the number of Wnt5b-positive signaling protrusions was significantly reduced (Fig. 5F). These data indicate that Gpc4 promotes the formation of signaling protrusions that deliver Wnt5b to neighboring cells, representing a novel mechanism that might account for the long-range functions of Gpc4.

### Endodermal Gpc4-labeled signaling filopodia contribute to the rescue of mesodermal and ectodermal defects

We sought to determine whether signaling filopodia contribute to the rescue of mesoderm and ectoderm by endodermally expressed GFP-Gpc4 in *gpc4* mutants. The small Rho GTPase Cdc42 is a well-known promoter of filopodia formation *in vitro* (Kozma, et al., 1995; Nobes and Hall, 1995). Recent studies in zebrafish showed that interference with Cdc42 activity by overexpression of dominant-negative (dn) Cdc42 (Cdc42T17N) prevents the formation of signaling filopodia *in vivo* (Cayuso, et al., 2016; Stanganello, et al., 2015). We tested whether such interference with Cdc42 activity could suppress the rescue of body length defects in the context of endodermal expression of Gpc4. We injected embryos with a small dose of *cdc42T17N* RNA (120 pg) and found that such injection had little impact on either the *gpc4*^*-/-*^ embryos or their siblings; posterior body length was comparable among the uninjected and injected embryos (Fig. 6A-D, I). Notably, in the *Tg*(*sox17:GFP-gpc4*) background, overexpression of Cdc42T17N also did not alter posterior body length in control siblings (Fig. 6G vs E, I), yet it led to a significant decrease in length in *gpc4*^*-/-*^ embryos (Fig. 6H vs F, I). Thus, inhibition of Cdc42 activity can partially suppress the rescue by endodermal expression of GFP-Gpc4.

**Figure 6.**
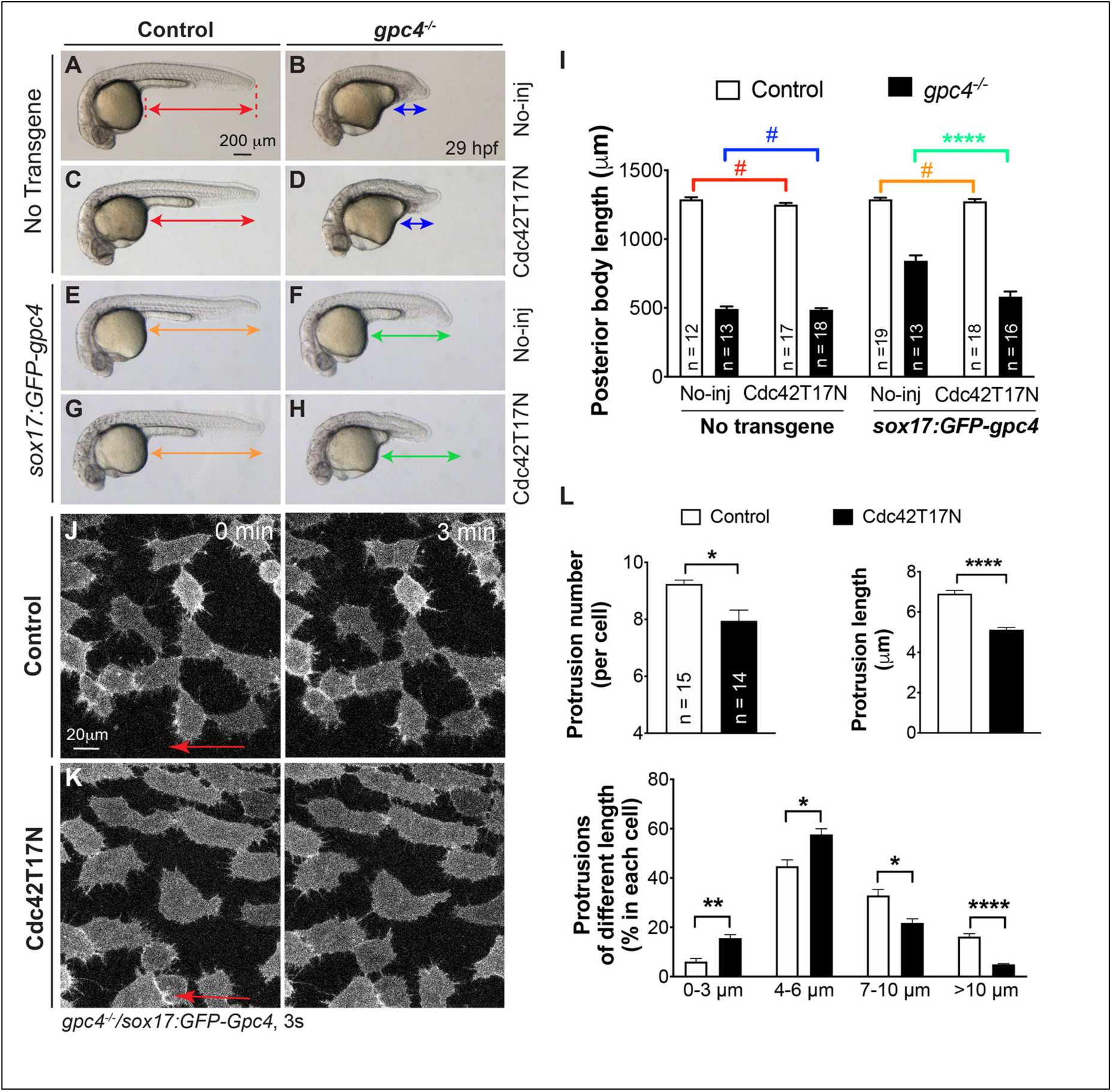
Suppression of GFP-Gpc4-labeled protrusions by dnCdc42 partially blocks rescue by endodermally expressed Gpc4. (A-H) Bright-field images of the indicated embryos. Lines with double arrows, length of the posterior body; lines of the same color are equal in length. (I) Average of the posterior body length in embryos shown in (A-H). The number of embryos tested in each group is indicated in the bar. Colored P values correspond to differences between the embryos in which the posterior body is marked with lines of the same color. (J-K) Snapshots from confocal time-lapse imaging performed on *gpc4*^*-/-*^*/Tg*(*sox17:GFP-gpc4*) embryos injected with *cdc42T17N* RNA and in uninjected controls (Movie 3). Red arrows, direction of migration of the endodermal cells. (L) The total number of protrusions in each endodermal cell, and protrusion length as well as the percentages of protrusions of different lengths, grouped into in 3 μm bins. The analyses were performed at 3-min intervals throughout the imaging sequences in control embryos (555 protrusions, 15 cells, 5 embryos) and embryos injected with *cdc42T17N* RNA (445 protrusions, 14 cells, 5 embryos). #, P>0.05, *, P<0.05, **, P<0.01, ****, P<0.0001; Student’s t-test.

To determine whether suppression by Cdc42T17N expression was due to its effects on GFP-Gpc4-labeled filopodia. We performed confocal time-lapse experiments on *gpc4*^*-/-*^ /*Tg*(*sox17:GFP-Gpc4*) embryos injected with the Cdc42T17N RNA. Endodermal cells of uninjected embryos extended many GFP-Gpc4-labeled, finger-like filopodia. In embryos injected with the *cdc42T17N* RNA, these cells formed fewer filopodia and those that did form were significantly shorter (Fig. 6K vs J, Movie 3). Quantification of the cell protrusions in the time-lapse movies (9 min) confirmed that the expresssion of Cdc42T17N reduced the total number, as well as the average length, of protrusions produced by each cell (Fig. 6L). Notably, the proportion of long protrusions (7-10 μm and >10 μm) was significantly decreased in Cdc42T17N-expressing endodermal cells (Fig. 6L). Thus, rescue of the body length in *gpc4*^*-/-*^ embryos by endodermal expression of GFP-Gpc4 was suppressed by the expression of Cdc42T17N, likely due to the suppression of filopodia formation.

To further assess the role of filopodia in the observed rescue, we used Latrunculin B (Lat B), a well-characterized inhibitor of actin polymerization. To avoid the effects of this drug on early development, starting at 80% epiboly, we treated embryos with 0.15 μg/ml Lat B, a dose that did not cause visible defects at 29 hpf in *gpc4* mutant and control sibling embryos (Fig. S7A-D). However, like injection with the Cdc42T17N RNA, the application of Lat B to *gpc4*^*-/-*^ /*Tg*(*sox17:*GFP-Gpc4) embryos significantly suppressed the rescue of trunk length (Fig. S7H vs F), although it also caused mild reduction of trunk length in the sibling embryos (Fig. S7G vs E). To test whether the suppression was due to the inhibition of protrusions, we treated *gpc4*^*-/-*^*/Tg*(*sox17:*GFP-Gpc4) embryos with Lat B from 80% epiboly to 2s. Confocal time-lapse experiments at 4s showed that Lat B treatment reduced both the number and length of endodermal GFP-Gpc4 positive protrusions (Fig. S7J-M, Movie 4). Notably, the proportion of long protrusions (longer than >7μm) was significantly reduced, whereas that of short protrusions was increased (Fig. S7N). Thus, some protrusions extended but failed to reach the neighboring cells and connect with them (Movie 4). Collectively, our data suggest that filopodia from GFP-Gpc4 expressing endodermal cells are critical for the observed rescue effects.

## Discussion

Our study leads us to propose a model whereby Gpc4 elicits its non-cell autonomous functions by regulating the formation of signaling filopodia (Fig. 7). In *gpc4*^*-/-*^ embryos, endodermal Gpc4-GFP expressing signaling filopodia transport Wnt proteins to neighboring tissues to rescue mesodermal C&E defects; when the filopodia were blocked, the resuce effects were suppressed.

**Figure 7.**
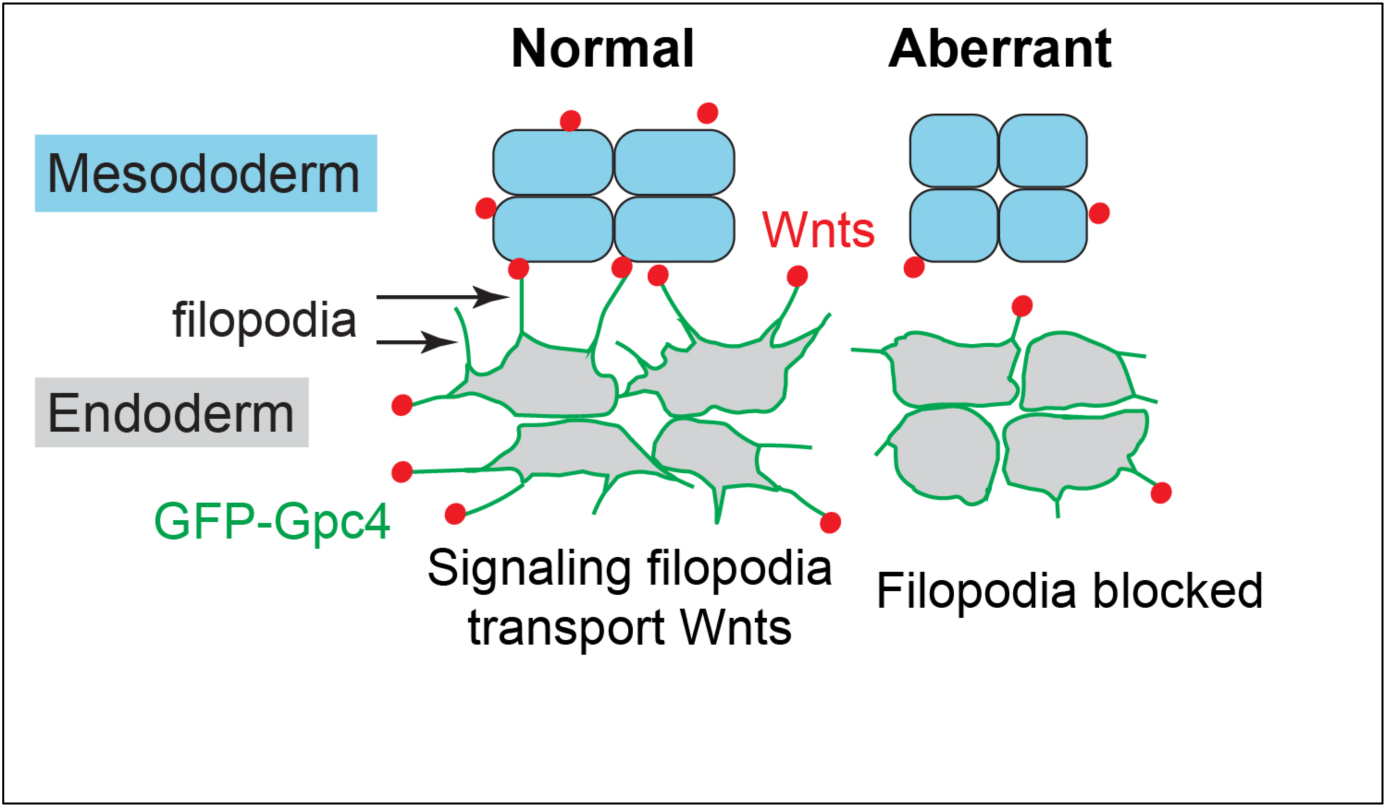
Model for how endodermal Gpc4-labeled signaling filopodia transport Wnt ligands to the mesoderm and ectoderm. In *gpc4*^*-/-*^ embryos, the signaling filopodia on endodermal cells deliver Wnts to *gpc4*-deficient mesodermal cells, producing cells of the shape necessary to restore body length. When filopodia formation is blocked, Wnt delivery is compromised and rescue of mesodermal C&E defects is suppressed.

### Endodermal expression of GFP-Gpc4 partially rescues the C&E defects of other germ layers

We provide multiple lines of evidence showing that in *gpc4*^*-/-*^ embryos, transgenic endodermal expression of GFP-Gpc4 not only completely rescues C&E defects in endoderm, but also partially rescues those in the mesoderm and ectoderm, indicating that Gpc4 functions both cell- and non-cell autonomously and that Gpc4 acts at long range. Furthermore, this rescue stems from the endoderm, as depletion of the endoderm abolishes such rescue. Explanations that would account for the partial rescue of mesoderm and ectoderm include the following: the level of Gpc4 expression driven by the *sox17* promoter is much lower than that driven by the endogenous *gpc4* promoter; and the expression of Gpc4 in the endoderm is not sufficient to compensate for a lack of Gpc4 expression in other layers of *gpc4*^*-/-*^ embryos.

Gpc proteins influence signaling pathways by interacting with signaling molecules. In zebrafish, Gpc4 cooperate with Wnt11f2 to regulate mesodermal C&E (Topczewski, et al., 2001), but it is not clear how it does this. Our study shows that Gpc4 can physically bind both Wnt5b and Wnt11f2, and that inhibiting the expression of both Wnt5b and Wnt11f2 at doses that do not cause significant embryogenic defects in body length can suppress Gpc4-mediated rescue of the mesoderm in *gpc4*^*-/-*^ embryos. These data suggest that in *gpc4*^*-/-*^ embryos the local concentrations of these Wnt proteins in the mesoderm are altered and can be partially restored by endodermal Gpc4. However, our study does not provide information on the distribution of these endogenous ligands *in vivo*; this will require knock-in reporter lines for Wnt5b and Wnt11f2. It is also possible that Wnt proteins cannot function properly without Gpc4 and that the endodermal Gpc4 compensates for loss in these germ layers. It would be interesting to test if Gpc4 expression in the mesoderm can rescue the endoderm defects in *gpc4*^*-/-*^ embryos. Future studies will also investigate whether and how Gpc4 regulates the distributions and functions of Wnt proteins to influence mesoderm C&E.

HS chains are thought to be critical for the interactions between Gpc proteins and morphogens (Yan and Lin, 2009; Hacker, et al., 2005). However, we found that Gpc4ΔGAG, which lacks the HS chain attachment site, is expressed on the cell membrane and binds both Wnt5b and Wnt11f2. Thus, the GAG domain of Gpc4 is not the sole region that binds Wnt and it is possible that it does not bind Wnt at all. In fact, the core Gpc protein has been shown to bind morphogens as well (McGough, et al., 2020; Kirkpatrick, et al., 2006; Capurro, et al., 2005). Nevertheless, HS chains can enhance the ability of Gpcs to bind signaling molecules (Yan, et al., 2009) or directly recruit morphogen-transporting vesicles (Eugster, et al., 2007). Although the CRD domain of GPC3 can bind Wnt3a (Li, et al., 2019), our zebrafish Gpc4 construct lacking this domain was not expressed on the plasma membrane (Fig. 3H). This prevents us from pursuing the functional significance of CRD in zebrafish Gpc4, and the exact region of Gpc4 that bind Wnt5b and Wnt11f2 remains to be determined.

### GPI cleavage is not necessary for the non-cell autonomous functions of Gpc4

Gpcs function both cell- and non-cell autonomously. However, little is known about how they achieve their non-cell autonomous functions. Our finding that endodermal Gpc4 expression rescues the C&E defects in other germ layers provides an unprecedented opportunity to study the role of Gpc4 in communication among tissues.

The GPI anchor of a Gpc can be cleaved from the plasma membrane to generate a soluble form of the protein that is transported to other sites and elicits its function at a distance (Hacker, et al., 2005; Lin, 2004). Here, we identified AA517-557 of Gpc4 as the potential GPI region (Fig. 3). Thus, it is possible that endodermal Gpc4 is cleaved and transported to other germ layers. However, our surprising discovery that transgenic expression of a membrane-bound form of Gpc4 in the endoderm is able to rescue defects in the mesoderm of *gpc4* mutants suggests that membrane localization, rather than cleavage of the GPI anchor, is necessary for its function in other germ layers. Furthermore, the fact that Gpc4 without the GPI anchor (Gpc4Δ517-557) failed to rescue the Gpc4 mutant and its overexpression instead causes C&E defects (data not shown) indicates that Gpc4 lacking the GPI membrane anchor does not function properly. We speculate that the presence of this form of the protein in the extracellular matrix interferes with the function of endogenous Gpc4 or Wnt/PCP signaling. However, we cannot rule out the possibility that regions of Gpc4 other than the GPI anchor can be cleaved. Thus, it remains to be determined whether zebrafish Gpc4 can be cleaved by certain enzymes at specific regions.

### Endodermal Gpc4-labeled signaling filopodia could be responsible for mesodermal C&E rescue

Accumulating evidence shows that ligand gradients allow physical communication among tissues and directly deliver signaling molecules including Wnt (Gonzalez-Mendez, et al., 2019; Stanganello and Scholpp, 2016; Kornberg and Roy, 2014). Notably, recent studies revealed that zebrafish blastula cells extend signaling filopodia that deliver Wnt8a to the neighboring cells, activating Wnt signaling (Stanganello, et al., 2015). Our study extends these findings and demonstrates that, in addition to blastula cells, endodermal cells extend signaling filopodia. Notably, our *in vivo* imaging shows that Gpc4 localizes to signaling filopodia that can bind mCerry-Wnt5b. However, given that Gpc4 bound both Wnt5b and Wnt11f2 *in vitro*, and inhibiting the expression of both proteins suppressed Gpc4-mediated rescue of the mesoderm, it is possible that Gpc4-labeled filopodia transport Wnt11f2 *in vivo* as well.

The fact that Gpc4 is present on signaling filopodia suggests that this protein might contribute to the regulation of these structures, consistent with findings that Drosphila Dally and Dlp are required for the spreading of cytonemes that deliver Hh (Gonzalez-Mendez, et al., 2017). A recent study showed some Gpcs, including Dlp and GPC4, but not Dally serve as lipid moiety reservoirs to allow binding and solubilizing Wg, facilitating Wnt transport (McGough, et al., 2020). Here, we provide evidence to show additional role of Gpc4 in transporting Wnt proteins, by promoting formaiton of long and productive signaling filopodia. Thus, the reduction in the proportion of long filopodia could affect the efficiency of ligand delivery to distant sites. It is also possible that Gpc4 helps to stabilize or elongate the protrusions. Notably, in *gpc4*^*-/-*^ embryos, the number of mCherry-Wnt5b-bound filopodia is significantly reduced, probably due to the loss of binding to Gpc4. Wnt5b-bound signaling filopodia extended from endodermal cells and retracted back, in some cases bringing mCherry-Wnt5b into the cells. These behaviors of filopodia could contribute to regulating the local concentrations of morphogens, and perhaps gradients, in the tissues. It will be interesting to monitor the influence of Gpc4 on the distribution endogenous Wnt proteins in live embryos in future.

Our findings indicate that Gpc4 regulates the formation of long signaling filopodia, consistent with a recent report that Wnt/PCP signaling actived by the receptor tyrosine kinase Ror2 can induce the formation of filopodia in zebrafish blastula cells (Mattes, et al., 2018). Thus, Wnt/PCP signaling could be involved in the formation of signaling filopodia, and the underlying mechanisms could contribute to many other developmental processes regulated by Wnt/PCP signaling. It will also be important to determine whether signaling filopodia contribute to other events in morphogenesis during which Wnt/PCP signaling is needed.

Our data indicate that Wnt5b transportation by signaling filopodia is likely responsible for the observed rescue because the rescue is significantly blocked when filopodia formation is suppressed by either the expression of dnCdc42 expression or treatment with Lat B. Although the suppression was not complete, we surmise that this was because high levels of dnCdc42Gpc4 expression or Lat B treatment impaired mesodermal C&E (data not shown). Thus our findings reveal a novel mechanism whereby Gpc4 regulates signaling pathways at a distance *in vivo*, forming signaling filopodia to regulate signaling molecules. Nonetheless, Gpc4 might use additional mechanisms such as exosomes for long-distance signaling (Melo, et al., 2015).

In summary, our study indicates that endodermal Gpc4 can affect other germ layers because of the formation of signaling filopodia. This novel mechanism appears to be relevant for interactions between Gpc4 (and potentially other HSPGs) and Wnt, as well as other signaling molecules such as Bmp and FGF (Venero Galanternik, et al., 2016; Strate, et al., 2015).

## Materials and Methods

### Zebrafish strains and maintenance

Zebrafish were maintained as described previously (Xu, et al., 2011). Animal protocols were approved by the University of Iowa Animal Care and Use Committee. Embryos were obtained by natural spawning and staged either according to morphological criteria or hours post fertilization (hpf) at 28 or 32°C unless otherwise specified, as described previously (Kimmel, et al., 1995). The following zebrafish lines were used in this study: AB*/Tuebingen, *Tg*(*sox17:memCherry*) (*Ye, et al*., *2015*), *gpc4/knypek*^*fr6*^ (Topczewski, et al., 2001), *sox32/casanova*^*s4*^ (Kikuchi, et al., 2001), *wnt11f2/silberblick*^*tz216*^ (Heisenberg, et al., 2000), and *wnt5b/pipetail*^*ti265*^ (Hammerschmidt et al. 1996). The *gpc4* mutant was genotyped using primers targeting an intron region of *gpc4* as described (Hu, et al., 2018). For the genotyping of *sox32*^*s4*^, *wnt11f2*^*tz216*^, and *wnt5b*^*ti265*^ mutants, genomic DNAs were amplified using the following primers and PCR amplicons were digested using restriction enzymes for specific patterns. For *sox32*^*s4*^, the forward primer is 5’-TACATGCAAGAAGCAGAAAGACTACGGATCCAGG and the reverse primer is 5’-ATGTTGCCTCGAAGTGGTATGATGAAGAGTGGTT; and KpnI digestion produces a band at 271 bp from the wild-type genomic DNA and bands at 233 bp and 38 bp from the mutant genomic DNA. For *wnt11f2*^*tz216*^, the forward primer is 5’-TAGTATTTGGGTGATTCCATTAGG and the reverse prime is 5’-GTGGTTGAGGCTTTACCTGTCT; FokI digestion produces bands at 403 bp and 134 bp from the wild-type DNA and a 537 bp band from the mutant DNA. For *wnt5b*^*ti265*^, the forward primer is 5’-GTCTCTGGGCACCCAAGGCCGCCTATGC and the reverse primer is 5’-CAAACTGGTCTACGAGTGACGTGCAGCGTTTGCTC; XbaI digestion produces a single band at 185 bp from wild-type DNAs and bands at 147bp and 38bp from mutant DNAs.

### Sequence alignment

Alignment of C-terminal amino acids of Gpc4 from mouse (ENSMUST00000033450.2), rat (ENSRNOT00000003282.5), human (ENST00000370828.3), *Xenopus* (ENSXETT00000011898.2) and zebrafish (ENSDART00000026569.8) was performed in Clustal X, a multiple sequence alignment program (Thompson, et al., 1997).

### Plasmid constructs

The following truncated Gpc4 constructs were generated by PCR: GFP-Gpc4-GPI (Δ24-516) in which AA24-516 are deleted so that only the N-terminal signal peptide (AA1-23) and C-terminal GPI attachment signal (AA517-557) of Gpc4 are included; GFP-Gpc4ΔGAG (Δ484-504), in which the HS chain attachment domain (GAG domain, AA484-504) is deleted; and GFP-Gpc4ΔCRD (Δ196-351), in which the potential CRD domain (AA196-351) is deleted. In these PCR reactions, pCS2dest GFP-Gpc4, in which EGFP is inserted after N-terminal signal peptide of Gpc4 (Hu, et al., 2018) (as the DNA template), and specific primers containing BstbI and XhoI restriction enzyme sites were used with Q5 high-fidelity DNA polymerase (New England Biolabs). The resulting amplicons were cut using BstbI and XhoI, and were then cloned into the pCS2dest GFP-Gpc4 plasmid cut by BstbI and XhoI. The pCS2dest GFP-Gpc4Δ517-557-Sdc4TM construct contains Gpc4 AA1-517, the transmembrane domain (T144-173) and a partial intracellular sequence (R174-L185) of zebrafish Syndecan4 (Sdc4) (NM_001048149.1). The coding region of AA144-185 s*dc4* was amplified from cDNAs obtained from 18s-zebrafish embryos using the following primers: 5’-CTAT<u>ACCTGGT</u>ACAGAAGTGCTTGCAGCTGTT-3’ and 5’-TATA<u>CTCGAG</u>TTACAGGTCGTAACTTCCTTCGTCT-3’ (the underlining indicates the SexAI and XhoI restriction sites, respectively). Because the conserved cytosolic domain (G186-A201) of Sdc4 binds to intracellular signaling molecules (Multhaupt, et al., 2009) and we sought to prevent such interactions, we removed this domain and included only twelve amino acids of the cytosolic sequence (R174-L185) in this construct. The amplicon was digested with SexAI and XhoI, and was then cloned into the pCS2dest GFP-Gpc4 plasmid following its digestion with SexAI and XhoI. pCS2 Flag-Gpc4 was a gift from Dr. Jacek Topczewski (Northwestern University), and the Flag epitope was inserted after the N-terminal signal peptide of Gpc4. pCS2 Flag-Gpc4ΔGAG (Δ484-504) was generated by PCR as described above and cloned into the pCS2 Flag-Gpc4 plasmid following its cleavage by BstbI and XbaI. pCS2 lifeact-RFP was generated by amplifying the coding sequence of lifeact-RFP using Abp140-17aaRuby-nos1-3’UTR (Kardash, et al., 2010) as a template and cloning the product into the pCS2 vector.

### Generation of transgenic lines

*Tg*(*sox17:memGFP/H2A-mCherry*), *Tg*(*sox17:GFP-gpc4*) and *Tg*(*sox17:GFP-Gpc4*Δ*517-557-sdc4TM*) were generated using a Tol2-based Multi-Site Gateway system (Invitrogen, Carlsbad, CA) (Kwan, et al., 2007; Villefranc, et al., 2007). *pME*-GFP-Gpc4 was a gift from Dr. Topczewski (Northwestern University). The *pME-*GFP-Gpc4Δ517-557-TM was generated by amplifying the coding sequence of GFP-Gpc4Δ517-557-TM from pCS2dest GFP-Gpc4Δ517-557-TM plasmid (see above) using primers containing attB sites, and the resulting PCR products were recombined into a pDONR221 vector using BP Clonase II Enzyme mix (Invitrogen, Carlsbad, CA). A 5’-entry vector containing a *sox17* promoter was used to express genes specifically in the endoderm (Woo, et al., 2012). A 3’-entry vector *p3E*-polyA and the destination vector pDestTol2pA2 were used for Multi-Site Gateway cloning. Transgenic lines were established using standard techniques as described previously (Xu, et al., 2014). The founders were bred to generate multiple stable lines. We utilized lines that express GFP-Gpc4 at a modest level and in which embryogenesis was normal. For genotyping of *Tg*(*sox17:GFP-gpc4*), the following primers were used to generate a amplicon at 241 bp: forward primer (5’-TGTTTACAGTATGTATGTCTGTGGTGG), which targets the N-terminal signal peptide of Gpc4; reverse primer (5’-GTCAGGGTGGTCACGAGGG), which targets the open frame sequences of GFP.

### RNA expression and Morpholino (MO) injection

mRNA and MOs were injected at the one-cell stage at the doses indicated. Capped messenger RNAs were synthesized using the mMessage mMachine kit (Ambion, Foster City, CA) and were injected into one-cell embryos. RNAs encoding the following genes were used: *memCherry* (75 pg), *H*_*2*_*A-mCherry* (75 pg), lifeact-RFP (200 pg), *mCherry-wnt5b* (125 pg) (Lin, et al., 2010), *cdc42T17N* (Nobes and Hall, 1995) (120 pg), *GFP-gpc4*, and *GFP-gpc4* truncated constructs (200 pg each). Previously validated morpholino antisense oligonucleotides (MOs) targeting the following genes were used: *sox32* (4 ng, 5’-CAGGGAGCATCCGGTCGAGATACAT) (Wong, et al., 2012); *wnt5b* (1 ng, 5’-GCAAACACAATAATTTCTTACCACC) (Cirone, et al., 2008); *wnt11f2* (5 ng, 5’-ACTCCAGTGAAGTTTTTCCACAACG) (Muyskens and Kimmel, 2007); *p53* (1.5 ng, 5’-GCGCCATTGCTTTGCAAGAATTG) (Robu, et al., 2007). All MOs were co-injected with the *p53* MO to inhibit potential p53-dependent cell death induced by MO off-targeting effects (Robu, et al., 2007).

### Whole-mount *in situ* hybridization (WISH)

Digoxigenin-labeled antisense RNA probes targeting *hgg1, dlx3, shh* (Marlow, et al., 1998), *foxa3* (Odenthal and Nusslein-Volhard, 1998), *deltaC* (Haddon, et al., 1998), *krox-20* (Oxtoby and Jowett, 1993), and *tbxta* (*ntl*) (Schulte-Merker, et al., 1994) were synthesized by *in vitro* transcription. WISH was performed as previously described (Thisse and Thisse, 2008; Lin, et al., 2005). For double ISH, a probe against *hgg1* was labeled with fluorescein (Roche) and detected by INT/BCIP (Roche) staining.

### Latrunculin B treatment

Embryos at 80% epiboly were dechorionated in glass dishes and treated with Lat B (428020, Sigma-Aldrich). This inhibitor of actin polymerization was applied at a dose of 0.15 μg/ml (diluted in 0.3% Danieau buffer with 1% DMSO). Control embryos were treated with 0.3% Danieau buffer containing 1% DMSO. Embryos were treated from 80% epiboly to 29 hpf at 28°C and then subjected to bright-field imaging. For the confocal time-lapse experiment, embryos were treated from 80% epiboly to 2s at 28°C, and then washed thoroughly with 0.3% Danieau buffer before mounting for imaging.

### Cell transfection, immunoprecipitation and Western blotting

HEK293 cells were transiently transfected with pCS2 Myc-Wnt5b (Lin, et al., 2010), pCS2 Myc-Wnt11f2 (both were gifts from Diane Slusarski), pCS2 Flag-Gpc4 (a gift from Jacek Topczewski), pCS2 Flag-Gpc4ΔGAG and pCDNA Flag-Jnk* (MKK7B2Jnk1a1) (Young, et al., 2014) (a gift from Ray Dunn), as described previously (Sun, et al., 2011). Thirty-six hours after transfection, the cells were collected in lysis buffer [150 mM NaCI, 50 mM Tris-HCI (pH 7.5), 5 mM EDTA, 0.5% NP-40, 50 mM NaF] (Ohkawara, et al., 2003) containing protease inhibitors and subjected to sonication using an ultrasonic processor (GE505, Sonics & Materials) operated at 5 cycles of 1s on, 10s off. For immunoprecipitation, the cell lysates were incubated with Myc monoclonal antibody (9E10) (MA-1-980, ThermoFisher Scientific, Waltham, MA) coupled to Protein G magnetic beads (88848, ThermoFisher Scientific, Waltham, MA) overnight at 4°C. The precipitates were collected using a magnetic separation rack (CS15000, ThermoFisher Scientific, Waltham, MA) and washed with lysis buffer six times. Protein samples were resolved by SDS-PAGE and examined by Western blotting using an Amersham imager 600 detection system (GE Healthcare). The following antibodies were used for immunoblotting: anti-Flag M2 (1:2000, F3165, Sigma-Aldrich), anti-c-Myc (9E10) (1:2000, MA1-980, ThermoFisher Scientific), goat anti-mouse IgG, and light chain-specific HRP conjugate (1:2000, 115-035-174, Jackson ImmunoResearch Laboratories).

### Microscopy and image analysis

For still imaging, fixed embryos were mounted in 80% glycerol and live embryos were mounted in 2.5% methylcellulose. Still epifluorescence, WISH and bright-field images were taken as described previously (Hu, et al., 2018).Confocal images for the *in vivo* protein localization assay and fixed sample were taken on a laser-scanning confocal inverted microscope (Zeiss LSM880, Carl Zeiss, Inc.) with EC Plan-Neo 40x/NA 1.3 oil or LD C-Apo 40×/NA 1.1 water objectives. Z-stacks were acquired at optimal intervals using the following settings: 1024×1024 pixel, 9 speed, 4 averaging.

For confocal time-lapse imaging, embryos were embedded in 0.7% (for embryos aged to less than 10 hpf) or 1% (for embryos aged beyond 10 hpf) low melting-point agarose using glass-bottom dishes, and images were taken at 28°C using Zeiss LSM880 with a LD C-Apo 40×/NA 1.1 water objective and a temperature-controlled stage. Endoderm cells in the posterior region of embryos were focused for imaging cell protrusions. For moive 1, z-stacks of 15 μm were acquired at 1.5 μm intervals every 15 seconds using the following settings: 512×512pixels, 8 speed, 2 averaging. For other time-lapse imaging, z-stacks of 13.5 μm were acquired at 1.5 μm intervals every 30 seconds using the following settings: 1024×1024 pixels, 9 speed, 4 averaging.

Images of the same type were acquired using the same settings, and all images were processed using Fiji software, edited, and compiled using Adobe Photoshop® and Adobe Illustrator software. To evaluate the anterior-posterior axis, we focused on the posterior region of *gpc4* mutant embryos, which show obvious defects. We used an easy and accurate method to quantify the posterior body length, tracing from the starting point of the yolk extension to the tip of the tail using the segmented-line or straight-line tool in Fiji software.

Cell protrusions were assessed by taking snapshots from the confocal time-lapse movies every 3 minutes and determining the average number and length of cell protrusions in each cell. To assess the number of GFP-Gpc4 expressing protrusions that bind Wnt5b-mCherry, we analyzed the confocal time-lapase movies performed on *Tg*(*sox17:GFP-gpc4*) injected with mCherry-Wnt5b RNA. For each cell, the number of protrosions labeled with Wnt5b-mCherry were manually tracked from 9-minutes movies at 30-seconds intervals with seven z-planes. The length of cell protrusion was measured from the starting point on cell membrane to the tip of the protrusion using a straight-line tool in Fiji software.

### Statistical analysis

Data were compiled from two to three independent experiments and are presented as the mean ± SEM. Statistical analyses were performed in GraphPad Prism (GraphPad Software) using unpaired two-tailed Student’s *t*-tests with unequal variance, and P<0.05 was considered significant. The number of cells and embryos analyzed in each experiment is indicated in the figure legends.

## Acknowledgments

This work was supported by funding to FL from the National Science Foundation (IOS-1354457) and NIH (1R56DK123610-01). We are grateful to: Jacek Topczewski (Northwestern University) for technical suggestions and providing the plasmid constructs; Diane Slusarski (University of Iowa), Bruce Appel (University of Colorado), Ray Dunn (Institute of Medical Biology, Singapore) and Erez Raz (University of Münster) for providing the plasmid constructs. We thank other members in the laboratories of Drs. Lin and Robert Cornell (University of Iowa), as well as Lilianna Solnica-Krezel and Diane Sepich (Washington University in St. Louis), for their technical suggestions. We also thank Matthew Culver and Matthew Murry for excellent fish care and technical support.

## Author Contributions

B.H. and F.L. conceived the ideas and designed experiments; B.H., A.K.B., J.J.R., Y.Y.G., N.T.N., H.S., S.S and C.S. performed the experiments; B.H. and F.L. wrote the manuscript.

## Competing interests

The authors declare no competing or financial interests.

## Notes

### Competing Interest Statement

The authors have declared no competing interest.

