## Supplemental Figures 1-7 for "Glypican4 mediates Wnt transport between germ layers via signaling filopodia"

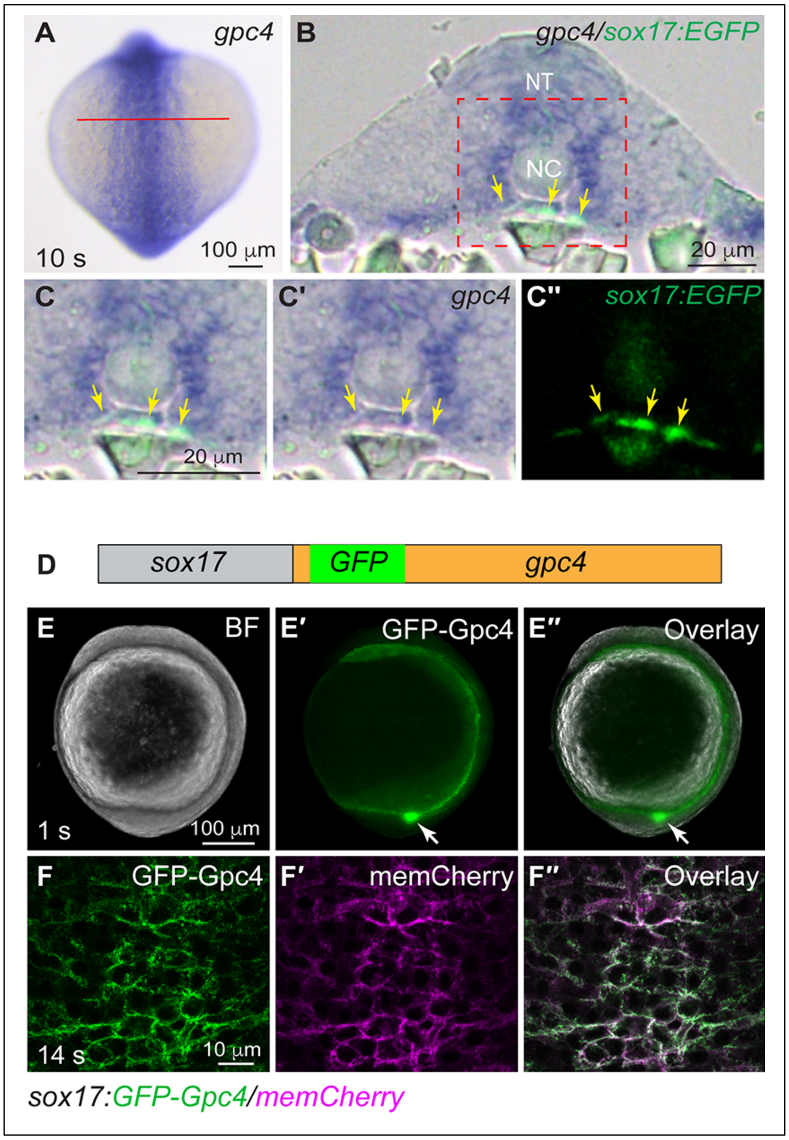


**Figure S1. *gpc4* is expressed in posterior endoderm and transgenic line that expresses Gpc4 in the endoderm.**

(A-C′′) Expression of *gpc4* transcript in the posterior region of a *Tg(sox17:EGFP)* embryo at 10s, as detected by whole-mount *in situ* hybridization (WISH). Posterior dorsal view, with anterior up. Red line, estimated plane for cross-sectioning. (B-C′′) Transverse sections of the embryo. (C-C′′) Higher magnification images of the region shown in red dashed box in (B). (C) Overlay of ISH panel (C′) and anti-GFP immunofluorescence staining panel (C′′). Yellow arrows: endoderm. NT: neural tube, NC: notochord. (D) Schematic depiction of transgene in Tg(*sox17*:GFP-Gpc4). GFP (green box) is inserted after the N-terminal signal peptide of Gpc4, with expression driven by the endoderm-specific promoter *sox17* (gray box). (E-E′′) Expression of transgenic GFP-Gpc4 at 1s. (E) Bright-field image. (E′) Epifluorescence image of GFP. (E′′) Overlay of (E) and (E′). White arrows: Kupffer's vesicle. (F-F′′) A representative confocal z-stack image at 14s, showing the expression of GFP-Gpc4 (F) and mCherry (F′) on the plasma membranes of endodermal cells, and (F′′) overlay of (F) and (F′).

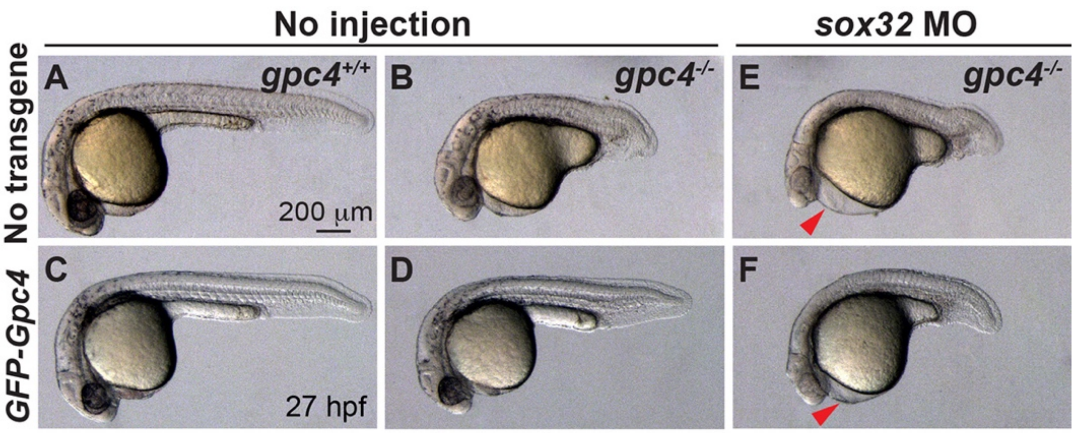


**Figure S2. Rescue of shortened body length in *gpc4* mutants by endodermal GFP-Gpc4 expression is suppressed by inhibition of endoderm formation.**

Bright-field images of embryos of the indicated genotypes obtained from incrossing *gpc4*$^{+/-}$ /*Tg(sox17:*GFP-Gpc4*)* fish, injected with a *sox32* MO or not (No injection). Red arrowheads: pericardial edema.

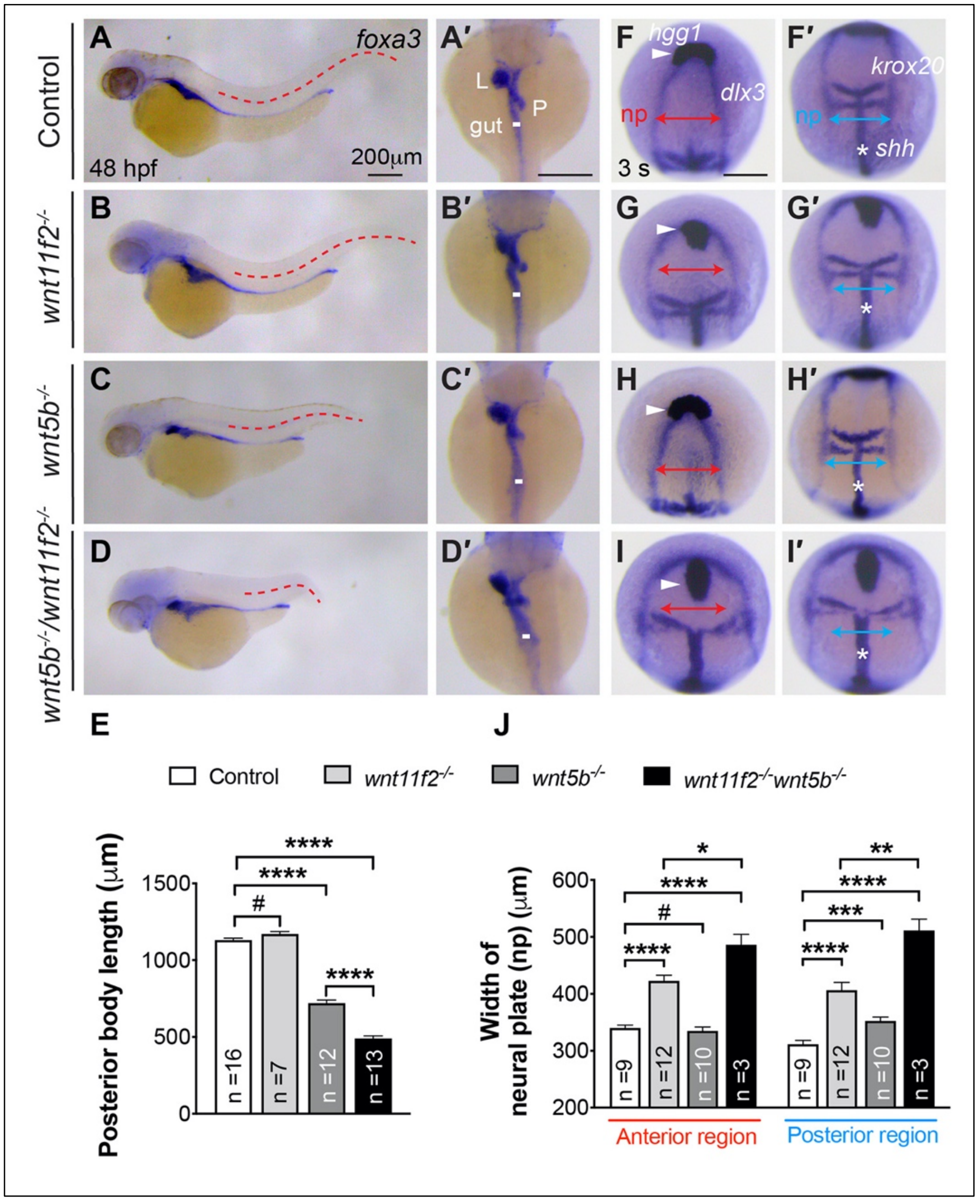


**Figure S3. Synergistic C&E defects in mesoderm and endoderm of *wnt5b* and *wnt11f2* mutants.** (A-D′) *foxa3* expression, as assessed by WISH, in the indicated embryos, showing morphology of the gut, liver (L), and pancreas (P). (A-D) Lateral view, (A′-D′) Dorsal view. Red dashed lines: length of posterior body; white lines, width of the gut-tube. All lines of the same type are equal in length. (E) Average posterior body length in embryos in (A-D). (F-I′) Expression of *hgg1* (white arrowheads), *dlx3* (np), *krox20* and *shh* (*), as detected by WISH in the indicated embryos. Red and blue lines with double arrows: width of neural plate (np) in the anterior and posterior region, respectively. All lines of the same color are equal in length. (J) Average width of the neural plate in the anterior and posterior regions in embryos shown in (F-I′). #, P>0.05, *, P<0.05, **, P<0.01, ***, P<0.001, ****, P<0.0001; Student's t-test. Scale bar: 200 μm.

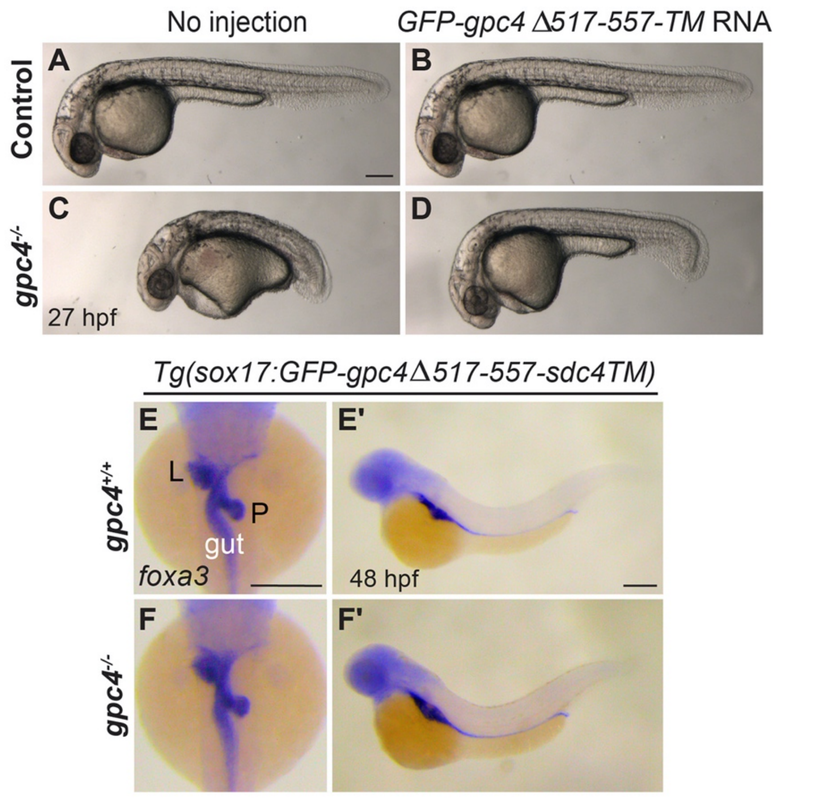


**Figure S4. GPI cleavage of Gpc4 does not drive mesoderm rescue.**
(A-D) Bright-field images of the indicated embryos at 27 hpf. (E-F′) Expression of *foxa3,* as detected by WISH, in the indicated embryos at 48 hpf, showing the morphology of the gut, liver (L), and pancreas (P). (E-F) Dorsal view. (E′-F′) Lateral view. Scale bars: 200 μm.

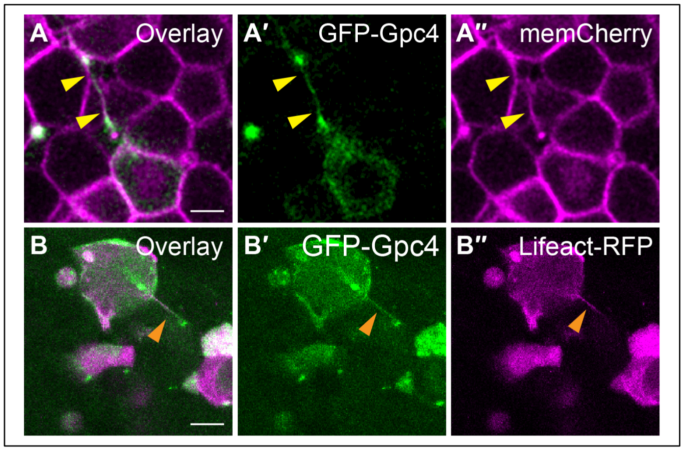


**Figure S5. Gpc4 contributes to the formation of actin-based filopodia.** (A-A′′) Live, confocal images focused on a zebrafish blastula cell (labeled with memCherry, in magenta, A′′) that expresses both GFP-Gpc4 (A′) and nuclear $H_2A$-mCherry (in magenta, A′′). Yellow arrowheads: GFP-positive prolonged protrusion. (B) Live, confocal images of blastula showing cells co-labeled with GFP-Gpc4 (B′) and Lifeact-RFP (in magenta, B′′). Orange arrowhead: protrusion co-labelled with GFP and Lifeact-RFP. Scale bar: 10μm.

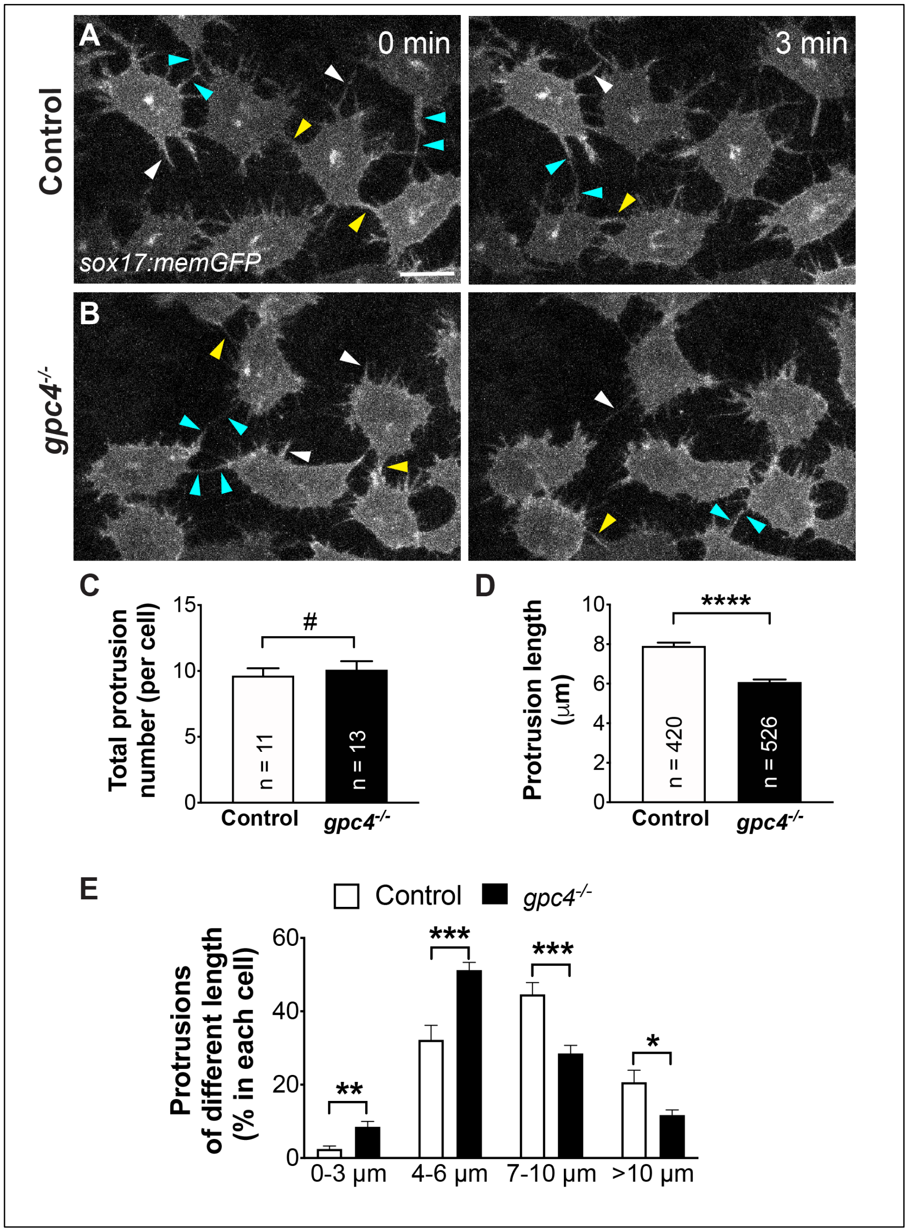


**Figure S6. Gpc4 is necessary for formation of long protrusions.**

(A-B) Snapshots from confocal time-lapse imaging performed on *gpc4*$^{-/-}$ */Tg(sox17:memGFP)* embryos and their siblings. White arrowheads, protrusions are present in the space between endodermal cells; yellow arrowheads, protrusions that connect with neighboring endodermal cells; cyan arrowheads, protrusions linking two cells. (C) Average of the total number of protrusions in each endodermal cell (at 3-min intervals throughout the imaging sequences) in *gpc4*$^{-/-}$ embryos (526 protrusions, 13 cells, 4 embryos) and their siblings (420 protrusions, 11 cells, 3 embryos). (D) Average length of protrusions in (C). (E) The percentages of protrusions in (D) of different lengths grouped into 3-μm bins. #, P>0.05, *, P<0.05, **, P<0.01, ***, P<0.001, ****, P<0.0001; Student's t-test.

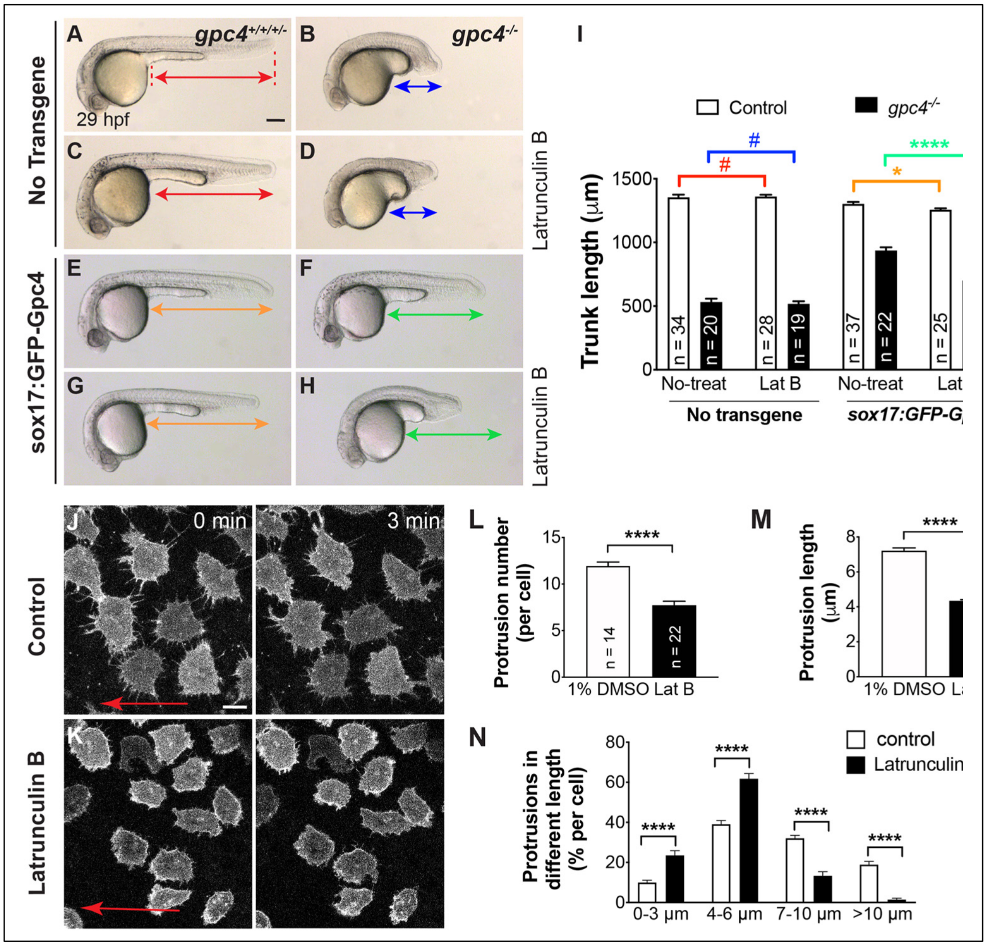


**Figure S7. Inhibition of actin polymerization by Latrunculin B blocks rescue by endodermal expression of Gpc4.**

(A-H) Bright-field images of the indicated embryos. Lines with double arrows, length of the posterior body; lines of the same color are equal in length. (I) Average of the posterior body length in embryos shown in (A-H). The number of embryos tested in each group is indicated in the bar. Colored P values correspond to the embryos in which the posterior body is marked with lines of the same color. (J-K) Snapshots from confocal time-lapse imaging performed on *gpc4*$^{-/-}$*/Tg(sox17:GFP-gpc4)* embryos treated with Latrunculin B and 1% DMSO (Movie 4). Red arrows, direction of migration of the endodermal cells. (L) Average of the total number of protrusions in each endodermal cell (at 3-min intervals throughout the imaging sequences) in embryos treated with 1% DMSO (673 protrusions, 14 cells, 3 embryos) and embryos treated with Latrunculin B (684 protrusions, 22 cells, 4 embryos). (M) Average length of protrusions in L. (N) The percentages of protrusions of different length in M, grouped into 3-µm bins. #, P>0.05, *, P<0.05, ****, P<0.0001; Student's t-test.

**Movie 1. GFP-Gpc4 expressing endodermal cells extend long, GFP-labeled cellular protrusions.**
Confocal time-lapse experiment was performed on *Tg(sox17:GFP-gpc4)* embryo at 4s, using a Zeiss LSM880 confocal microscope with a LD C-Apo 40×/NA 1.1 water objective. Images were acquired at 15-sec intervals and the movie plays at 5 frames/sec.

**Movie 2. Protrusions labeled by memGFP in *gpc4* mutant and its sibling.**
Confocal time-lapse experiment was performed on 3s *Tg(sox17:memGFP)* embryo, using a Zeiss LSM880 confocal microscope with a LD C-Apo 40×/NA 1.1 water objective. Images were acquired at 30-sec intervals and the movie plays at 5 frames/sec.

**Movie 3. GFP-Gpc4-labeled protrusions in *gpc4* mutant embryos are suppressed by Cdc42T17N.**
Confocal time-lapse experiments were performed on *gpc4* mutant in *Tg(sox17:GFP-gpc4)* embryos at 3s, using a Zeiss LSM880 confocal microscope with a LD C-Apo 40×/NA 1.1 water objective. Images were acquired at 30-sec intervals and the movie plays at 5 frames/sec.

**Movie 4. GFP-Gpc4-labeled protrusions in *gpc4* mutant embryos are suppressed by Latrunculin B.**
Confocal time-lapse experiments were performed on *gpc4* mutant in *Tg(sox17:GFP-gpc4)* embryos at 3s, using a Zeiss LSM880 confocal microscope with a LD C-Apo 40×/NA 1.1 water objective. Images were acquired at 30-sec intervals and the movie plays at 5 frames/sec.
